# EDC-3 and EDC-4 Regulate Embryonic mRNA Clearance and Biomolecular Condensate Specialization

**DOI:** 10.1101/2024.03.04.583404

**Authors:** Elva Vidya, Yasaman Jami-Alahmadi, Adarsh K. Mayank, Javeria Rizwan, Jia Ming Stella Xu, Tianhao Cheng, Rania Leventis, Nahum Sonenberg, James A. Wohlschlegel, Maria Vera, Thomas F. Duchaine

## Abstract

Animal development is dictated by the selective and timely decay of mRNAs in developmental transitions, but the impact of mRNA decapping scaffold proteins in development is unknown. This study unveils the roles and interactions of the DCAP-2 decapping scaffolds EDC-3 and EDC-4 in the embryonic development of *C. elegans*. EDC-3 facilitates the timely removal of specific embryonic mRNAs, including *cgh-1, car-1,* and *ifet-1* by reducing their expression, and preventing excessive accumulation of DCAP-2 condensates in somatic cells. We further uncover a novel role for EDC-3 in defining the boundaries between P-bodies, germ granules, and stress granules. Lastly, we show that EDC-4 counteracts EDC-3 and engenders the assembly of DCAP-2 with the GID (CTLH) complex, a ubiquitin ligase involved in maternal-to-zygotic transition (MZT). Our findings support a model wherein multiple RNA decay mechanisms temporally partake in the clearance of maternal and zygotic mRNAs throughout embryonic development.

## Introduction

Regulated messenger RNA (mRNA) decay is critical for developmental processes. For instance, the majority of maternally inherited transcripts are degraded in early embryos of all animal species before maternal-to-zygotic transition (MZT)^1,2^. Failure to do so leads to embryonic lethality and impaired establishment of the soma-germline boundaries in *C. elegans*^3,4^. The timely and specific decay of mRNAs in this transition is dictated by a compendium of trans-acting factors that include RNA-binding proteins and the ∼22-nt long non-coding microRNAs (miRNAs)^5,6^. For example, *let-7* miRNAs direct the decay of *lin-41* mRNA during L4-to-adult transition in *C. elegans*^7^.

The 5’-end of mRNAs is modified with a 7-methylguanosine (m7G) “cap” linked to the first transcribed nucleotide via a 5’-5’ triphosphate linkage^8^. Hydrolysis of the 5’ mRNA cap (herein referred to as Decapping) is a critical step in mRNA decay in development and homeostatic pathways^9^. mRNA decapping is catalyzed by a decapping holoenzyme consisting of the catalytic enzyme Dcp2 (*C. elegans* DCAP-2) and its main cofactor Dcp1 (DCAP-1)^10,11^. The decapping holoenzyme physically associates with and is activated by scaffolding proteins, among which EDC-3 (**<u>E</u>**nhancer of **<u>D</u>**e**<u>C</u>**apping) and EDC-4 have been best characterized in yeast and metazoan cell cultures, respectively^11–13^. Notably, EDC-3 has a well-documented role in promoting phase separation of a decapping complex in yeast^14–17^ but is dispensable for P-bodies formation in metazoan models^18–21^, while EDC-4 is the primary scaffold for DCAP-2 accumulation in P-bodies^22,23^.

Additional factors such as LSm14 (CAR-1, for **<u>C</u>**ytokinesis, **<u>A</u>**poptosis, **<u>R</u>**NA-associated**<u>-1</u>**), Ddx6 (CGH-1, for **<u>C</u>**onserved **<u>G</u>**ermline **<u>H</u>**elicase**<u>-1</u>**), 4E-T (IFET-1, for e**<u>IF</u>**4**<u>E</u> <u>T</u>**ransporter**<u>-1</u>**) and PatL1 (PATR-1, for **<u>PAT</u>**1 **<u>R</u>**elated**<u>-1</u>**) are linked to decapping enhancement by promoting RNA binding activity, by stabilizing the active conformation of Dcp1/2, and by facilitating the formation of higher-order decapping assemblies^24–30^. In contrast, in *Drosophila* early embryos and *C. elegans* oogenesis, a complex formed of CAR-1, CGH-1, and IFET-1 represses translation and prevents mRNA decapping and decay^31,32^, suggesting molecular coordination between the two related processes.

All known decapping scaffolding proteins concentrate in cytoplasmic biomolecular condensates called P-bodies, along with the 5’-3’ exonuclease XRN-1, a fraction of the miRNA-induced silencing complex, the CCR4-NOT deadenylase complex, and RNA-binding proteins^14,33–35^. In *C. elegans* embryos, P-bodies are found in the somatic and germline blastomeres and are distinct from P/germ granules, which specifically form in the germline lineage^36,37^. Originally proposed to be active sites for mRNA decay^38^, several lines of evidence support the idea that P-bodies instead store and protect silent mRNAs from decapping and decay^33,39–41^. However, as P-bodies seem to form as a consequence of translational repression, some papers even challenged whether they represent active regulatory sites *per se*^21,42^. As the support for these differing models stems from distinct species and biological contexts, it is possible that the role(s) of P-bodies in the fate of mRNAs depends on their composition, which diverges with cell types and developmental stages^43^. Thus, understanding how decapping proteins are scaffolded and functionally organized across development will provide insights into the functions of P-bodies.

Despite accumulating evidence for biochemical interactions between decapping scaffolds across various species and contexts^44^, their functional relationship *in vivo* is not fully understood. Notably, it is currently unclear whether and how EDC-3 and EDC-4 functionally intersect during development, and whether they function redundantly or independently of each other. Furthermore, immunofluorescence on *C. elegans* embryos revealed that IFET-1, CAR-1, and CGH-1 proteins are dramatically reduced in somatic blastomeres starting from the 4-cell stage while they remain expressed in the germline blastomeres in subsequent stages^45–47^. This suggests that the decapping machineries, as well as well as their biomolecular condensate assemblies, may be subject to specialization and spatiotemporal regulation during *C. elegans* development.

Here, we document the functions of EDC-3 and EDC-4 in scaffolding DCAP-2 in cytoplasmic foci across *C. elegans* early embryonic development. We uncover a specific role for EDC-3 in the timely degradation of *cgh-1, car-1,* and *ifet-1* mRNAs in the developing embryos and an antagonistic function for EDC-4. We further reveal a novel function for EDC-3 in determining the boundaries across co-existing biomolecular condensates.

## Results

### Opposing functions for EDC-3 and EDC-4 in DCAP-2 foci *in vivo*

To trace the expression and localization of DCAP-2 during *C. elegans* embryonic development, we employed CRISPR-Cas9^48,49^ to introduce a GFP linked to the C-terminal end of *dcap-2*. No phenotype observed in *dcap-2* null animals (*dcap-2(ok2023)*) was visible in the *dcap-2::GFP* strain, suggesting that DCAP-2 remains functional^50^. DCAP-2 was then followed throughout embryonic development, grouping embryos as 1∼2-cell, 4∼6-cell, 7∼16-cell, ∼24-cell, ∼50-cell and ∼100-cell for analysis **(Figure S1A)**. A gradual increase in DCAP-2 expression was readily noticeable throughout development, with P-bodies being visible starting at the 4-cell stage **(Figures 1A-B)**. For clarity and considering our focus on DCAP-2 as the main marker for P-bodies, herein we will refer to DCAP-2-containing P-body-related condensates as DCAP-2 foci. To quantify and characterize DCAP-2 foci, we re-purposed StarDist^51,52^, a deep-learning-based segmentation plugin in Fiji^53^, to automate the detection and quantification of DCAP-2 foci’s number, diameter and intensity **(Figure S1B)**. Within our imaging parameters, wild-type (WT) embryos accumulated on average ∼250 DCAP-2 foci between 4∼6-cell stage and up to ∼1,000 foci at ∼100-cell stage **(Figure 1C)**. The intensity of DCAP-2 foci also significantly increased, by 48% between 4∼6-cell and ∼24-cell embryo populations, after which their intensity remained stable **(Figure 1D)**. DCAP-2 foci were on average the largest (∼554 nm in diameter) in 7∼16-cell embryos and then slightly shrank after ∼24-cell (∼536 nm diameter) **(Figure 1E)**. Thus, DCAP-2 expression gradually increases during embryonic development and forms increasingly brighter foci starting at the 4-cell stage, which coincides with the onset of zygotic transcription^54^.

**Figure 1.**
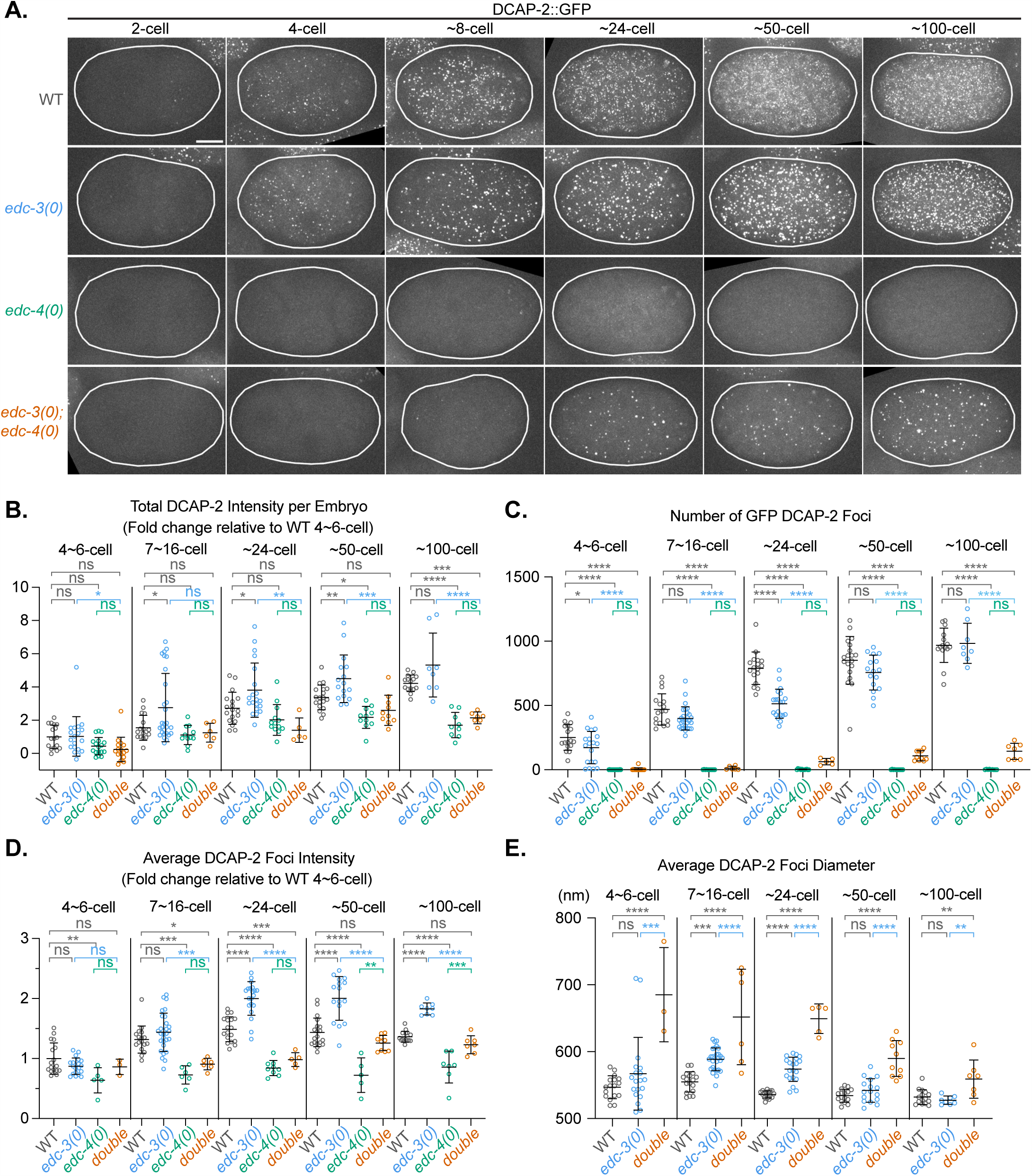
Opposing functions for EDC-3 and EDC-4 in DCAP-2 foci *in vivo*. (A) Representative photomicrographs of live embryos expressing DCAP-2::GFP. Scale bar is 10 µm. Corresponding brightfield images are shown in **Figure S1A**. (B) Total DCAP-2 intensity per embryo. Represented as fold change of the mean intensity of all slices (sum projection), relative to WT at 4∼6-cell. (C) Number of DCAP-2 foci per embryo as detected with StarDist. Examples of foci detection are shown in **Figure S1B**. (D) Average DCAP-2 foci intensity per embryo. Represented as fold change of the mean intensity of maximum z-projection from all slices, relative to WT at 4∼6-cell. (E) Average of DCAP-2 foci’s Feret’s diameter per embryo. Results are representative of three technical and biological replicates wherein all genotypes were imaged in parallel. For (B)-(E), each data point represents one embryo and is summarized as mean ± standard deviation. Tukey’s multiple-comparisons test was conducted for each stage bin. P-values: (*) =0.01-0.05, (**) =0.001-0.01, (***) =0.0001-0.001, (****) =<0.0001, p≥0.05 are insignificant (ns) See also **Figure S1**.

EDC-3 and EDC-4 are two decapping scaffolds that physically interact with the DCAP-1/2 holoenzyme in metazoans and are thought to regulate its activity^12,24^. To investigate whether EDC-3 and EDC-4 function in parallel or independently of one another *in vivo*, we examined the impact of their individual and combined genetic depletions on DCAP-2 in *C. elegans* embryos. To interrogate the functions of EDC-3, we utilized the *edc-3(ok1427)* allele (herein referred to as *edc-3(0)*) which harbours a ∼1 kb deletion spanning six exons on *edc-3* **(Figure S1C)**^55,56^. We further generated an *edc-4(qe49)* allele (herein referred to as *edc-4(0)*) in which a premature stop codon was introduced in the middle of the second exon using CRISPR-Cas9 genome editing approach **(Figure S1D)**. We also generated rabbit polyclonal antibodies to detect EDC-3 and EDC-4 proteins and confirmed their depletion in *edc-3(0)* and *edc-4(0),* respectively **(Figure S1E-G)**.

Loss of EDC-3 led to unanticipated changes to DCAP-2 foci **(Figure 1A)**. On average, *edc-3(0)* accumulated fewer DCAP-2 foci than WT. The most striking difference occurred in ∼24-cell embryos wherein *edc-3(0)* formed on average 25% less DCAP-2 foci than WT **(Figure 1C)**. In addition, DCAP-2 foci were 35% brighter in *edc-3(0)* than in WT in ∼24, ∼50, and ∼100-cell embryos **(Figure 1D)**. DCAP-2 foci were also larger in diameter in *edc-3(0)* compared to WT, by 6% (30 nm) and 1.5% (8 nm), in 7∼16-cell and ∼24-cell embryos, respectively **(Figure 1E)**. Total DCAP-2 expression was also increased by 1.4-fold in *edc-3(0)* in comparison to WT, between 7∼16 and ∼50-cell embryos **(Figures 1A-B)**, a finding that was confirmed by western blot **(Figure 3G**, compare WT and *edc-3(0)*, quantified in **Figure 3H**). The observation that loss of *edc-3* concentrates DCAP-2 into fewer foci of greater brightness and size suggests that EDC-3 opposes DCAP-2 localization to foci.

In contrast to *edc-3(0)*, loss of *edc-4* led to ∼50% less total DCAP-2 than WT **(Figures 1A-B, 3G-H)**. *edc-4(0)* embryos accumulated nearly no detectable DCAP-2 foci across all stages **(Figures 1B-C)**. Thus, in contrast to EDC-3, EDC-4 stabilizes DCAP-2 and promotes its accumulation into foci in *C. elegans* embryo.

Strikingly, *edc-3(0)* rescued DCAP-2 foci formation in *edc-4(0)* **(Figure 1A)**. While the total DCAP-2 signal in *edc-3(0); edc-4(0)* embryos was comparable to *edc-4(0)* across all stages **(Figure 1B)**, 60 to 140 DCAP-2 foci were observed in *edc-3(0); edc-4(0)*, starting from ∼24-cell embryos **(Figure 1C)**. These foci were initially less intense (in ∼24-cell embryos) than WT DCAP-2 foci, by 51%, but became as bright as WT later in development **(Figure 1D)**. Notably, DCAP-2 foci in *edc-3(0); edc-4(0)* animals were larger in diameter than WT by 21% (113 nm), 10% (56 nm), and 5% (27 nm) in ∼24, ∼50, and ∼100-cell embryo cohorts, respectively. *edc-3(0); edc-4(0)*’s DCAP-2 foci were also larger on average than *edc-3(0)* by 13% (76 nm), 9% (48 nm), and 6% (32 nm) at corresponding stages. Together, our findings show that EDC-3 inhibits, while EDC-4 promotes DCAP-2 accumulation to cytoplasmic foci. Opposing functions of EDC-3 and EDC-4 maintain the accumulation and distribution of DCAP-2, and its foci association.

### EDC-3 and EDC-4 define the composition of DCAP-2 interactome

Our observation that EDC-3 inhibits DCAP-2 foci accumulation is in contrast with reports that EDC-3 promotes DCAP-2 foci accumulation in yeast^16,17,57^. We hypothesized that in *edc-3(0)* embryos, the fraction of DCAP-2 bound to EDC-3 is now liberated to interact with other co-expressed condensate scaffolds. Since EDC-4 is a primary driver for DCAP-2 foci accumulation **(Figure 1)**, we reasoned that a comparative proteomic analysis of DCAP-2 between WT and *edc-3(0); edc-4(0)* may reveal the alternative DCAP-2 scaffold(s) that compete with the two major scaffolds (EDC-3 and EDC-4). We thus immunoprecipitated (IP) a CRISPR-engineered 3XFLAG-tagged of DCAP-2 from WT or *edc-3(0); edc-4(0)* embryonic lysates and analyzed the eluates by liquid chromatography coupled to tandem mass spectrometry (LC-MS/MS)^58^, and determined the enrichment of co-IP proteins over the negative control (N2(WT) strain expressing untagged DCAP-2).

A total of 136 proteins were significantly enriched by IP of DCAP-2 in WT (p-adjusted<0.05, log2 fold change (FC)>1.8) **(Figure S2A, Table S1)**. Across 3 biological replicates, EDC-3 and EDC-4 were the top two interactors of DCAP-2 (log2FC EDC-3=9.45, EDC-4=12.81), along with XRN-1 (log2FC=7.32). DCAP-2 also interacted with known decapping scaffolds (DCAP-1, PATR-1, CGH-1, CAR-1, IFET-1), the CCR4-NOT deadenylase complex (NTL-1/LET-711, CCR-4, CCF-1, NTL-2, NTL-3, NTL-9, NTL-11), and other known P-body localized proteins (such as MEG-1^37^ and NHL-2^59^). We also detected novel interactions with DCAP-2, notably with all 5 subunits of the GID (CTLH) ubiquitin ligase complex, including the core scaffold GID-1 (RanBP9/10), its partner GID-8, the enzymatic subunits GID-2 (RMND5A) and MAEA-1, the oligomerization-promoting subunit GID-7 (WDR26), as well as an additional co-factor B0546.4 (YPEL5) (Discussed below)^60^.

Interestingly, more than 4-fold more proteins (a total of 551) were enriched by DCAP-2 IP in the *edc-3(0); edc-4(0)* genotype **(Figure S2B, Table S1)**. Most notably, DCAP-2 interactions with IFET-1, CAR-1, and CGH-1 were increased by 1.6, 2.0 and 1.8-fold, respectively, in *edc-3(0); edc-4(0)* embryos in comparison to WT **(Figure 2A)**. This is intriguing because IFET-1, CAR-1, and CGH-1 are normally enriched in the germ cells and are absent from somatic cells^45–47^, where DCAP-2 is abundant **(Figure 1)**. Furthermore, 9 germ-granule enriched proteins, specifically MEG-3, MEX-1, MEX-3, PGL-3, GLH-1, GLH-2, PRG-1, CSR-1, and RDE-12 did not interact with DCAP-2 in WT but became enriched in *edc-3(0); edc-4(0)* embryos (log2FC= 4.87, 4.37, 4.07 for MEG-3, MEX-1, and MEX-3, and ∼2.2 for the others) **(Figure 2B)**. Furthermore, the two primary markers for stress granules, GTBP-1 (G3BP1) and TIAR-1 (TIA1), as well as four EIF-3 subunits (EIF-3.B, C, E, K), became significantly enriched in DCAP-2 IP in *edc-3(0); edc-4(0)* embryos (log2FC = 3.54 and 3.15 for GTBP-1 and TIAR-1, 1.9-2.6 for the EIF-3 subunits). Of note, DCAP-2 interaction with XRN-1 was reduced by 2.6-fold in *edc-3(0); edc-4(0)* embryos compared to WT **(Figure 2A)**, but the interaction with DCAP-1, PATR-1, or the CCR4-NOT deadenylase complex remained unchanged (<1.2-fold difference between *edc-3(0); edc-4(0)* and WT) **(Figure 2C)**. Strikingly, in *edc-3(0); edc-4(0)* embryos, DCAP-2 lost its interactions with all subunits of the GID complex **(Figure 2A)**. In summary, loss of both EDC-3 and EDC-4 resulted in an overall reorganization of the DCAP-2 interactome; gaining interactions with not only CAR-1, CGH-1, and IFET-1, but also proteins from distinct biomolecular condensates (germ and stress granules), and impairing interactions with several other partners, such as the GID complex.

**Figure 2.**
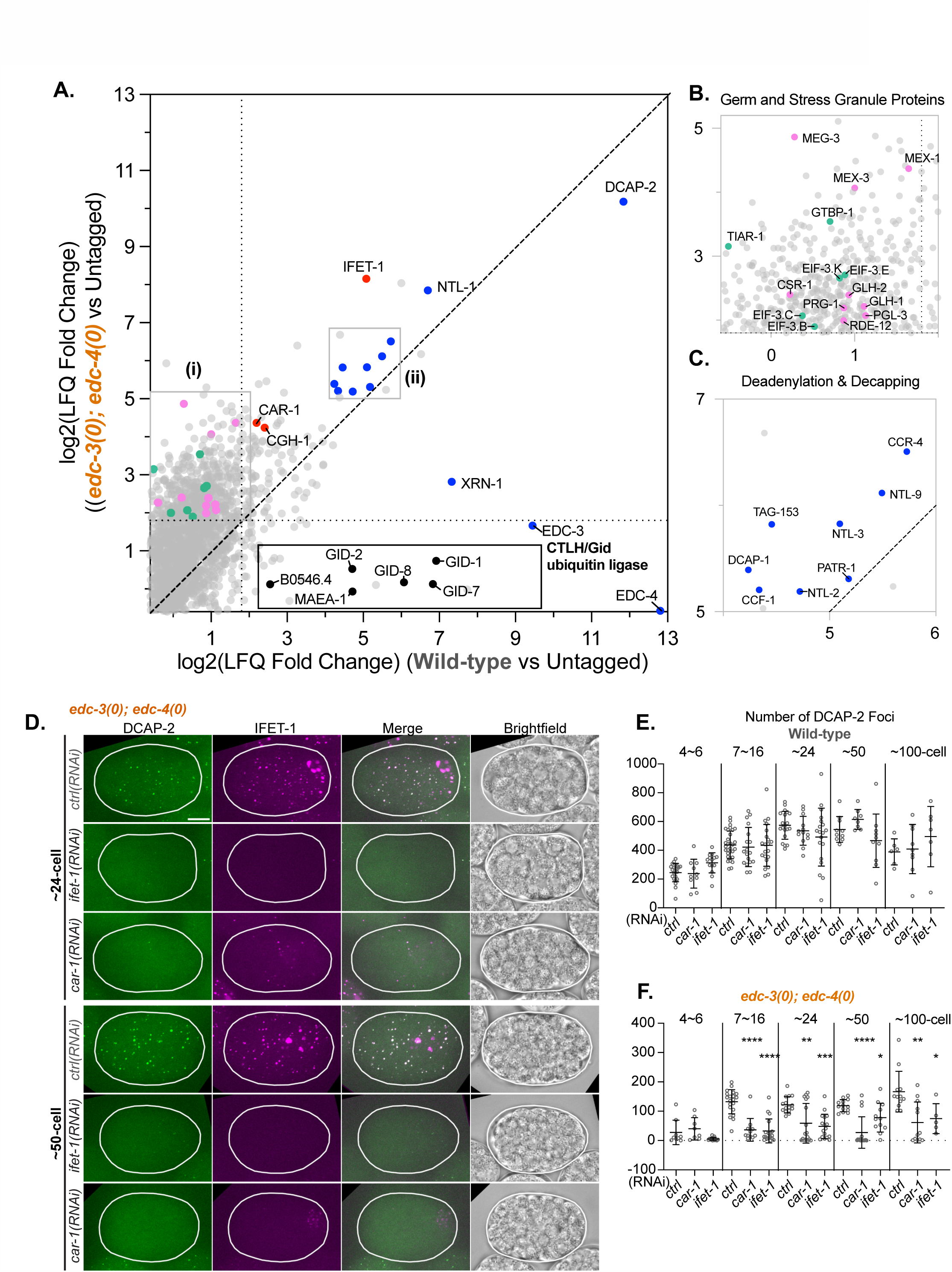
EDC-3 and EDC-4 define the composition of DCAP-2 interactome. (A) Log2(LFQ fold change) of proteins detected by LC-MS/MS of DCAP-2 immunoprecipitation (IP) from WT and *edc-3(0); edc-4(0)*. Log2(LFQ fold change) was computed using ProVision using the LFQ (Label Free Quantification) intensity of each protein in DCAP-2 IP compared to the untagged (N2) negative control (see Methods). Proteins of interest were coloured as follows: Blue = P-body-enriched decapping/decay and deadenylation proteins; Red = germline-enriched decapping scaffolds; Pink = Germ granule-enriched proteins; Green = Stress granule-enriched proteins; Black = GID complex subunits. Vertical and horizontal dotted lines indicate log2(LFQ Fold Change) cut-off of 1.8. The full list of proteins is listed in **Table S1**. (B) Proteins that did not significantly interact with DCAP-2 in WT and became significant interactors in *edc-3(0); edc-4(0)*. A zoom of box (i) from (A). (C) Zoom of box (ii) in Figure (A), highlighting P-body-localized proteins. (D) Representative photomicrographs of live ∼24-cell and ∼50-cell embryos expressing IFET-1::TagRFP and DCAP-2::GFP fed with double-stranded RNA targeting *ifet-1*, *car-1*, or a non-targeting control *(ctrl)* RNAi. Scale bar is 10 µm. (E) Number of DCAP-2::GFP foci in WT embryos as detected with StarDist. (F) Number of DCAP-2::GFP foci in *edc-3(0); edc-4(0)* embryos as detected with StarDist. (A-C) Results were obtained from three biological and technical IP replicates. (D-F) Results were representative of two biological and technical replicates wherein all genotypes were imaged in parallel. Dunnett test was conducted to determine the significance relative to *ctrl(RNAi)* in each stage bin. P-values: (*) =0.01-0.05, (**) =0.001-0.01, (***) =0.0001-0.001, (****) =<0.0001, p≥0.05 are insignificant and are not indicated on the graph. See also **Figure S2**.

To investigate which of the observed changes were driven by EDC-3 versus EDC-4, we surveyed DCAP-2 interactome in the *edc-3(0)*, and *edc-4(0)* single mutants. IP yielded a total of 374 significant interactors of DCAP-2 in *edc-3(0)* **(Figure S2C, Table S1)**, which included 3 out of the 9 germ granule proteins (MEG-3, GLH-2, and RDE-12) that were captured in *edc-3(0); edc-4(0)*, and 2 additional germ granule markers (DEPS-1 and LAF-1). GTBP-1 and 4 EIF-3 subunits (EIF-3.A, B, I and M) also interacted with DCAP-2 in *edc-3(0)*, but not TIAR-1. Furthermore, all GID complex subunits still interacted with DCAP-2 in *edc-3(0)*. In contrast, only 38 proteins were significantly enriched by DCAP-2 IP in *edc-4(0)* **(Figure S2D, Table S1)**. Importantly, neither the germ or stress granule-localized proteins (except for EIF-3.A), nor the GID complex interacted with DCAP-2 in *edc-4(0)* IP. DCAP-2 also lost its interactions with DCAP-1 and XRN-1 in *edc-4(0)*, but it maintained interactions with EDC-3, PATR-1, and most subunits of the CCR4-NOT deadenylase complex.

In summary, loss of EDC-3 alters the composition of DCAP-2 foci from normal P-bodies into foci that also accumulate germ granule-enriched and stress-granule-localized proteins. Meanwhile, EDC-4 promotes DCAP-2 interaction with the GID complex, as well as with its co-factor DCAP-1, and with XRN-1.

### The IFET-1/CAR-1/CGH-1 complex scaffolds DCAP-2 foci in the absence of EDC-3 and EDC-4

Since a complex formed by IFET-1, CAR-1, and CGH-1 orthologs scaffold P-bodies in other species^19,61–63^, we hypothesized that their increased interactions with DCAP-2 may lead to the formation of the foci observed in *edc-3(0); edc-4(0)* embryos **(Figure 1)**. Because complete knockout of *ifet-1*, *car-1*, or *cgh-1* resulted in sterile animals or embryonic lethal phenotypes^45–47,64^, we knocked down *ifet-1* or *car-1* by RNAi to assess their contribution for scaffolding DCAP-2 foci in WT or *edc-3(0); edc-4(0)* embryos **(Figure S2E**, knockdown efficiency of 81% and 64% for *ifet-1* and *car-1(RNAi)*, respectively). Neither *car-1* nor *ifet-1(RNAi)* altered DCAP-2 foci accumulation in WT embryos **(Figure 2E)**. In stark contrast, *car-1(RNAi)* or *ifet-1(RNAi)* in *edc-3(0); edc-4(0)* resulted in a 2 to 4.5-fold reduction in DCAP-2 foci in embryos of 7∼16 to ∼100-cell **(Figure 2D, F)**. Notably, 35-75% of *car-1(RNAi); edc-3(0); edc-4(0)* embryos formed less than 5 DCAP-2 foci in embryos staged between ∼24 and ∼100 cell. We also noted that *car-1(RNAi)* disassembled IFET-1 foci in *edc-3(0); edc-4(0)* embryos (**Figure 2D**). Together, our results demonstrate that IFET-1 and CAR-1, and likely CGH-1, scaffold DCAP-2 foci in the absence of EDC-3 and EDC-4.

### EDC-3 inhibits somatic accumulation of IFET-1, CAR-1, and CGH-1

Since the IFET-1, CAR-1, and CGH-1 complex is normally depleted from somatic cells^45–47^, and loss of EDC-3 leads to increased DCAP-2 association with germ cell-enriched proteins, we hypothesized that EDC-3 may suppress IFET-1, CAR-1, and CGH-1 in the soma. To test this, we examined IFET-1 expression in strains expressing a CRISPR-engineered IFET-1::TagRFP fusion and compared it to strains bearing individual or combined *edc-3(0)* and *edc-4(0)* alleles. In WT cohorts, IFET-1 was most highly expressed in 1∼2-cell embryos **(Figure S3E)**, and IFET-1 foci were sharply enriched in the germ cell lineage **(Figure S3A).** At 7∼16-cell, total IFET-1 expression decreased by 52% on average **(Figure S3E).** Somatic IFET-1 foci, which were smaller than the germ cell IFET-1 foci, were observed in the soma **(Figure S3B-C)**. WT ∼24-cell embryos accumulated on average 57 IFET-1 foci, of which only ∼34% were somatic **(Figures 3A, C, E)**. The remaining germ cell IFET-1 foci were perinuclear **(Figure 3A)**, consistent with characteristics of the recently coined germline P-bodies^37^. At ∼50-cell, germ cell IFET-1 foci continued to dim, mirroring total IFET-1 intensity, which was reduced by 63% in comparison to 4∼6-cell embryos **(Figures 3B, S3E)**. In ∼100-cell embryos, almost no IFET-1 foci could be detected under our imaging parameters and the total IFET-1 intensity remained very low **(Figures S3D-E)**. In contrast, *edc-3(0)* embryos accumulated on average 4.6 times more IFET-1 foci than WT at ∼24-cell (263 foci in *edc-3(0)* vs 57 foci in WT) **(Figures 3A, C)**, of which on average 83% were somatic **(Figure 4E)**. Later in development (∼50-cell to ∼100-cell embryo populations), *edc-3(0)* formed between 12 and 20-fold more IFET-1 foci than WT **(Figures 3B-C, S5D)**, and these foci were on average trending brighter than in WT (**Figure 3D)**. Importantly, somatic IFET-1 foci overlapped with DCAP-2 in *edc-3(0)* **(Figure 3A-B)**, in agreement with IFET-1 being a scaffold of DCAP-2 foci in the absence of EDC-3 **(Figure 2)**. In contrast, loss of *edc-4* did not result in a notable difference in IFET-1 total intensity, foci accumulation, and soma-germline distribution across all embryonic stages **(Figures 3A-D, S3A-E)**. Thus, EDC-3, but not EDC-4, suppresses the accumulation of somatic IFET-1 in DCAP-2 foci.

**Figure 3.**
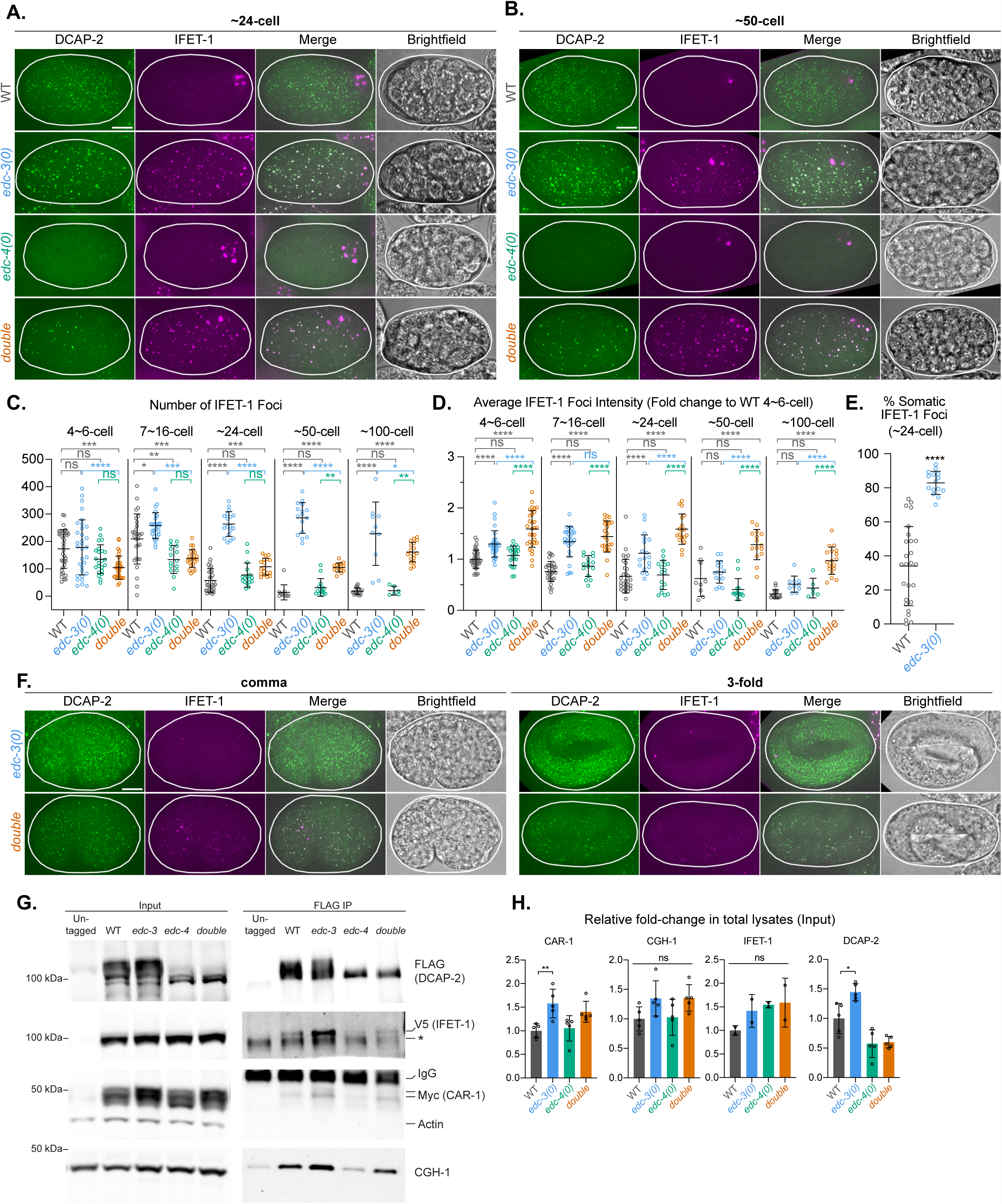
EDC-3 inhibits somatic accumulation of IFET-1, CAR-1, and CGH-1. (A, B, F) Representative photomicrographs of live (A) ∼24-cell; (B) ∼50-cell; (F) comma and 3-fold embryos expressing IFET-1::TagRFP and DCAP-2::GFP in WT, *edc-3(0)*, *edc-4(0)*, and *edc-3(0); edc-4(0)*. Scale bars are 10 µm. Additional stages are shown in **Figure S3A-D**. (C) Number of IFET-1::TagRFP foci per embryo as detected with StarDist. (D) Average IFET-1 foci intensity per embryo. Represented as fold change of the mean intensity of maximum z-projection from all slices, relative to WT at 4∼6-cell. (E) Percentage of somatic IFET-1 foci at ∼24-cell embryos in WT and *edc-3(0)*. Welch’s t-test was conducted to determine statistical significance. (****) indicates p-value <=0.0001. (G) Western blot from DCAP-2 FLAG co-immunoprecipitation experiment from mixed-stage embryos expressing *dcap-2::3XFLAG*; *ifet-1::V5*; *car-1::3XMyc* in WT, *edc-3(0)*, *edc-4(0)*, or *edc-3(0); edc-4(0)* double mutants. N2 strain (expressing untagged DCAP-2) serves as a negative control. 5% input (total lysates) was loaded. Actin served as a loading control in the input and negative control in the IPs. All images are representative of 3 technical and biological replicates. (H) Relative fold change of CAR-1, CGH-1, IFET-1, and DCAP-2 intensity from input lanes in (G), normalized to actin, relative to WT. (C), (D) and (H): Tukey’s multiple-comparisons test was conducted. p-values: (*) =0.01-0.05, (**) =0.001-0.01, (***) =0.0001-0.001, (****) =<0.0001, p≥0.05 are insignificant (ns). Data is summarized as mean ± standard deviation. See also **Figure S3**.

**Figure 4.**
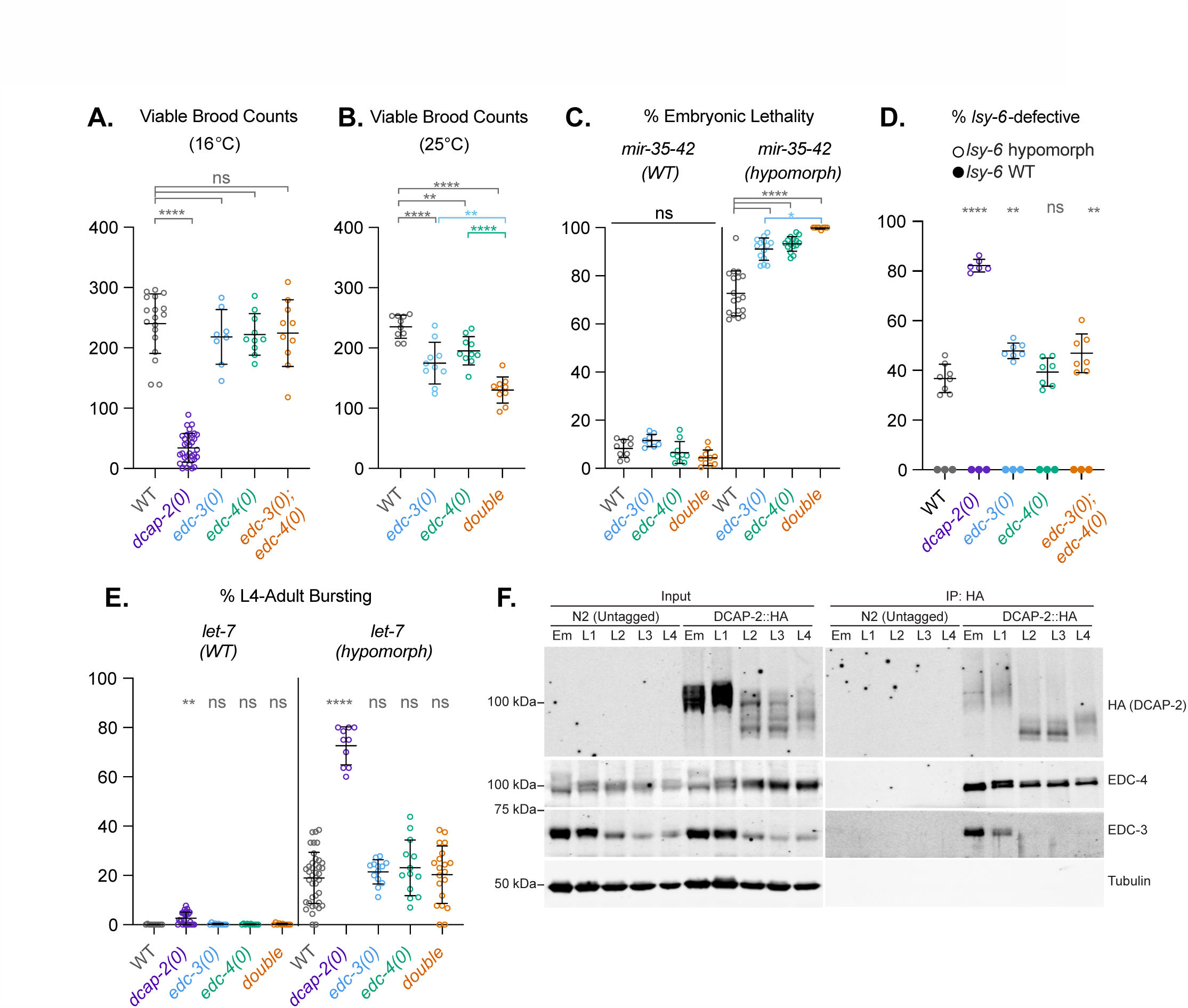
Selective roles for EDC-3 and EDC-4 in embryonic miRNA functions. (A-B) Viable brood counts in WT and decapping mutants at (A) 16°C and (B) 25°C. (C) Embryonic lethality in WT and decapping mutants with (Left) WT *mir-35-42* or (Right) hypomorphic for *mir-35-42* (i.e. *mir-35-41(nDf50)*). (D) Percentage of animals displaying defective silencing by the *lsy-6* miRNA, as marked by the presence of two, instead of one, *gcy-5p::GFP* ASER neurons. Filled data points at 0% were from WT *lsy-6* background. Clear data points were from the hypomorphic *lsy-6(ot150)* background. (E) Percentage of animals that died from vulval bursting in WT and decapping mutants in (Left) WT *let-7* or (Right) hypomorphic *let-7(n2853)* background. (A-E) Each data point represents counts from all F1s (for A-C and E) or at least 100 F1s (for D) from one hermaphrodite parent and summarized as mean ± standard deviation. For (A), (D), (E), Dunnett test was conducted to determine significance relative to WT. For (B) and (C), Tukey’s multiple comparison test was conducted. p-values: (*) =0.01-0.05, (**) =0.001-0.01, (***) =0.0001-0.001, (****) =<0.0001, p≥0.05 are insignificant (ns). (F) Western blots of DCAP-2, EDC-3, and EDC-4 proteins across embryonic (Em) and larval (L1-L4) stages of animals expressing DCAP-2::3XHA in (Left) total lysates and (Right) after IP with anti-HA beads. Tubulin served as loading control in the input and negative control in the IPs. Western blots are representative of two biological and technical replicates.

Interestingly, *edc-3(0); edc-4(0)* embryos also failed to clear somatic IFET-1 foci, but these foci were slightly different from those in *edc-3(0)*. Across all embryonic development, *edc-3(0); edc-4(0)* animals accumulated 30-60% less IFET-1 foci than in similarly staged *edc-3(0)* embryos, and these foci were brighter (by 30-50% in ∼24, ∼50, and 100-cell embryo cohorts) and 5-20% larger (30-120 nm in diameter) **(Figures 3A-D, S3F)**. Furthermore, at the post-differentiation stages, such as in the comma and 3-fold embryonic stages, IFET-1 foci were larger and brighter in *edc-3(0); edc-4(0)* embryos than in *edc-3(0)* **(Figure 3F)**. Since total IFET-1 intensity per embryo was comparable between *edc-3(0)* and the double *edc-3(0); edc-4(0)* mutant **(Figure S3E)**, we reason that loss of *edc-4* could further redistribute somatic IFET-1 in *edc-3(0)* into smaller but brighter foci.

Although we were not able to produce viable strains bearing fluorescently labelled CAR-1 and CGH-1 to assess their somatic distribution as with IFET-1, we were able to assess their expression using a CGH-1 polyclonal antibodies and a CRISPR-engineered Myc-tagged CAR-1, as well as their interaction with DCAP-2 in *edc-3(0)* and/or *edc-4(0)* mixed-stage embryonic lysates **(Figures 3G-H)**. CAR-1 and CGH-1 expression was elevated on average by 1.5 and 1.25-fold in *edc-3(0)* compared to WT **(Figures 3G** input lanes**, 3H)**, and loss of *edc-3* but not *edc-4* resulted in their increased co-IP with DCAP-2 **(Figure 3G** FLAG IP lanes). Of note, *edc-4*’s impact on the post-translational modification patterns of DCAP-2 will be described elsewhere (in preparation).

Taken together, our data support a model whereby EDC-3 suppresses the accumulation of the IFET-1/CAR-1/CGH-1 complex in the soma. While EDC-4 on its own does not affect the overall clearance of somatic IFET-1/CAR-1/CGH-1 complex, it affects IFET-1 localization into foci in the absence of EDC-3.

### Selective roles for EDC-3 and EDC-4 in embryonic miRNA functions

mRNA decapping and decay, along with translational repression, enact gene silencing pathways instigated by trans-acting factors, such as miRNAs^65,66^. To assess the importance of decapping scaffolds EDC-3 and EDC-4 *in vivo*, we investigated the impact of their loss on viability, and on hypomorphic alleles of three distinct miRNA families, namely *miR-35-42*, *lsy-6* and *let-7* miRNAs. When grown at 16°C, animals lacking *edc-3* and/or *edc-4* were viable and were as fertile as WT **(Figure 4A)**. However, when shifted to a higher temperature (25°C), *edc-3(0)* and *edc-4(0)* produced on average 26% and 17% less progeny than WT, respectively **(Figure 4B)**. Reduction in brood size was additive, as *edc-3(0); edc-4(0)* animals yielded broods on average 45% smaller than WT. This indicates that the pathways enacted by *edc-3* and *edc-4* are functionally disrupted under stress. In comparison, loss of DCAP-2 (*dcap-2(ok2023);* herein called *dcap-2(0)*) resulted in an 86% reduction in brood size compared to WT at 16°C **(Figure 4A)**, although this was primarily due to mothers being egg-laying defective (Egl)^50^.

Next, we assessed the impact of *edc-3(0)* and *edc-4(0)* on embryonic (*miR-35-42*, *lsy-6*) mRNA silencing pathways. Towards this, we crossed *edc-3(0)* and *edc-4(0)* with the *mir-35-41(nDf50)* strain, a hypomorphic allele of the *miR-35-42* family^67,68^. On its own, this hypomorphic allele exhibited ∼70% embryonic lethality at 16°C **(Figure 4C)**. *dcap-2(0)* was synthetic lethal with *mir-35-41(nDf50),* as we could not obtain a single viable homozygous *dcap-2(0); mir-35-41(nDf50)* animal. Individual *edc-3(0)* or *edc-4(0)* mutants exacerbated *mir-35-41(nDf50)* embryonic lethality to 91% and 93%, while their combined deletion exacerbated lethality to ∼99% **(Figure 4C)**. These results show that decapping is required for the function of *mir-35-42* and demonstrate that *edc-3* and *edc-4* facilitate silencing by this essential embryonic miRNA family. *edc-3(0)* and *edc-4(0)* were also crossed with a hypomorphic allele of *lsy-6(ot150)*^69^, a miRNA involved in left/right patterning of ASE chemosensory neurons (ASEL and ASER) through a decision that takes place in late (post-differentiation) embryonic stages, specifically between the bean and 3-fold stages^70,71^. The strain also carried an integrated GFP reporter driven by the ASER-specific *gcy-5* promoter^70^. On its own, 37% of *lsy-6(ot150)* animals were defective in silencing, as indicated by the appearance of two ASER neurons **(Figure 4D)**. In *dcap-2(0)*, this defect was exacerbated to 89%, indicating that decapping is important for robust silencing by *lsy-6*. In comparison, 48% and 47% of *edc-3(0)* and *edc-3(0); edc-4(0)* animals showed impaired *lsy-6* silencing **(Figure 4D)**, which marked a slight but significant increase over WT. In contrast, the *edc-4(0)* allele did not exacerbate the *lsy-6* phenotype, suggesting that it is dispensable for silencing by the *lsy-6* miRNA.

We further assessed *edc-3* and *edc-4*’s role in the larval functions of *let-7* miRNAs. The hypomorphic *let-7(n2853)* allele leads to 19% of animals bursting through the vulva at the L4-to-adult transition^72^ **(Figure 4E)**. The *dcap-2(0)* mutant on its own exhibited 2% of vulval bursting in the population and the same allele exacerbated the *let-7* hypomorphic phenotype up to 73%, highlighting the importance of the decapping enzyme for robust *let-7* function **(Figure 4E)**. Interestingly, and in contrast with embryonic miRNAs, neither individual nor combined *edc-3(0) and edc-4(0)* mutations had a significant effect on this function of *let-7* miRNAs **(Figure 4E)**.

A possible explanation for the selective involvement of EDC-3 and EDC-4 in the functions of embryonic miRNAs is that they may have restricted developmental expression domains. To begin assessing this possibility, we surveyed EDC-3 and EDC-4 expression and interaction with DCAP-2 in lysates isolated from across developmental time course **(Figure 4F)**. In western blots, EDC-3 was most highly expressed **(left panel)**, and specifically co-IP with DCAP-2 in embryos and at the L1 stage **(right panel).** In contrast, EDC-4 was ubiquitously expressed and co-IP with DCAP-2 in all embryonic and larval stages. Notably, while distinct isoforms or post-translationally modified forms of DCAP-2 could be observed across developmental stages, their interaction with EDC-4 was retained **(Figure 4E, right panel)**.

We thus conclude that EDC-3 and EDC-4 selectively partake in the functions of embryonic miRNAs. We note that while their distinct expression domains may account for some differences in physiological functions, they also highlight a possible involvement of other decapping scaffolds, for example post-embryonically. Furthermore, we cannot rule out functional compensation by other silencing mechanisms directed by miRNAs, such as translational repression (see Discussion).

### The GID complex is dispensable for somatic IFET-1 foci clearance

We next turned to investigate the mechanisms underlying EDC-3-mediated downregulation of the IFET-1/CAR-1/CGH-1 complex. Since the GID ubiquitin ligase complex promotes the degradation of the IFET-1/CAR-1/CGH-1 complex orthologs in *Drosophila*^73,74^ and all members of the GID complex were among the top interactors of DCAP-2 in our proteomic analysis **(Figure 2A)**, we hypothesized that the GID complex promotes the somatic degradation of IFET-1/CAR-1/CGH-1 in *C. elegans*. For this purpose, we generated a *gid-1(qe121)* allele (herein called *gid-1(0)*) harbouring a 42 bp insertion followed by a 126 bp deletion spanning the second to last exon-intron junction **(Figure 5A left panel)**. RT-qPCR revealed a >99% depletion of *gid-1* mRNA, confirming an efficient knockout **(Figure 5A right panel)**. We also generated a *gid-2(qe122)* allele (herein called *gid-2(mut)*) in which an essential and invariant cysteine residue in the RING domain was mutated to serine **(Figure 5B)**. Mutation of the equivalent cysteine in either *S. cerevisiae* or *X. laevis* ablated GID-2 ubiquitin ligase activity^75,76^.

**Figure 5.**
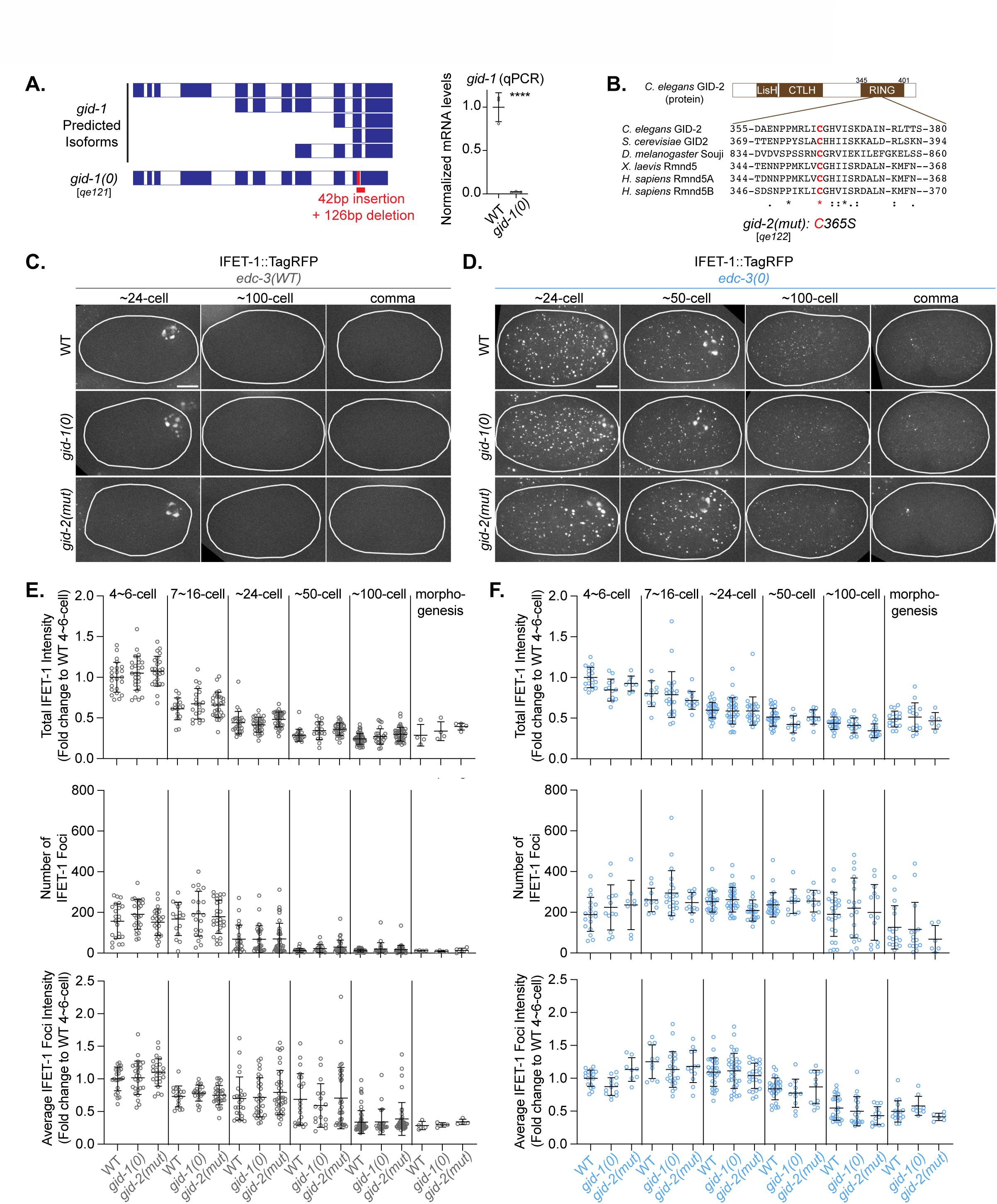
The GID complex is dispensable for somatic IFET-1 foci clearance. (A) Schematic representation of the *gid-1* genomic locus and null allele validation. (Left) *gid-1* locus and predicted isoforms where blue and white boxes indicate exons and introns, respectively. The location of genetic lesion *gid-1(qe121)* (i.e, *gid-1(0)*) is indicated. (Right) Relative expression of *gid-1* mRNA in *gid-1(0)* embryos as measured by RT-qPCR and normalized to *act-1* mRNA. Data is presented as mean ± standard deviation from three independent biological replicates. Welch’s t-test was conducted to determine statistical significance. (****) indicates *p*-value <=0.0001. (B) Schematic representation of GID-2 protein and conservation of the catalytic cysteine residue. GID-2 sequences from indicated species were aligned with Clustal Omega. Asterisk (*) indicates position with fully conserved residues, period (.) indicates conservation between weakly similar residues, and colon (:) indicates conservation between strongly similar residues. The *gid-2(qe122)* catalytic mutant (i.e. *gid-2(mut)*) bearing a C365S mutation (in red) was generated by CRISPR-Cas9. (C-D) Representative IFET-1::TagRFP photomicrographs of live embryos expressing IFET-1::TagRFP; DCAP-2::GFP in WT, *gid-1(0)*, or *gid-2(mut)* in (C) WT or (D) *edc-3(0)*. Scale bars are 10 µm. Corresponding brightfield and DCAP-2::GFP images are shown in **Figure S4A**. (E-F) Total IFET-1::TagRFP intensity, number of IFET-1 foci as detected with StarDist, and average IFET-1 foci intensity in (E) WT or (F) *edc-3(0*). To note, WT in (D) and (F) refers to *edc-3(0); gid-1(WT); gid-2(WT)*. (C-F) Results are representative of three technical and biological replicates wherein all genotypes were imaged in parallel. Each data point represents one embryo and data is summarized as mean ± standard deviation. Dunnet test was conducted to determine statistical significance relative to WT in each stage bin and differences were deemed insignificant (p≥0.05). See also Figure S4.

We reason that GID complex involvement could be reflected by the persistence of somatic IFET-1 foci past ∼24-cell in *gid-1(0)* or *gid-2(mut)*, mimicking the observation in *edc-3(0)* embryos. Instead, somatic IFET-1 foci were properly and timely cleared from either *gid-1(0)* and *gid-2(mut)* and were indiscernible from WT, including in ∼24-cell embryos and beyond **(Figure 5C)**. Similarly, total IFET-1 expression, the number of IFET-1 foci and the average IFET-1 foci intensity in *gid-1(0)* and *gid-2(mut)* were indistinguishable from WT across all embryonic stages **(Figure 5E)**. These data show that the GID complex is not responsible for the persistent somatic expression of IFET-1 or its foci observed in *edc-3(0)*. They also concur with the observation that DCAP-2 interaction with the GID complex is independent of EDC-3 **(Figure S3C).**

Since GID complex’s interaction with DCAP-2 is EDC-4-dependent **(Figure S2D)**, we next assessed the possibility that EDC-4 recruits the GID complex to mark somatic IFET-1 for degradation and that occurs in the absence of EDC-3. For this, IFET-1 foci persistence in *gid* mutants was profiled in *edc-3(0)* embryos, but IFET-1 total intensity, foci number and average foci intensity were indistinguishable from *edc-3(0)* alone **(Figures 5D, F)**. Lastly, we examined the possibility that the GID complex may instead affect DCAP-2 abundance or foci accumulation **(Figure S4)**. However, we did not observe any significant difference in total DCAP-2 intensity, in number of DCAP-2 foci, or in average foci intensity in *gid-1(0)* or *gid-2(mut)* in the presence or absence of *edc-3* **(Figure S4)**. Overall, and unlike in Drosophila^73,74^, these data do not support a critical role for the GID complex in clearing somatic IFET-1/CAR-1/CGH-1 complex or DCAP-2 foci from *C. elegans* embryos.

### EDC-3 promotes the somatic and germline clearance of *ifet-1*, *car-1*, and *cgh-1* mRNAs

We hypothesized that EDC-3 promotes the clearance of somatic *ifet-1*, *car-1,* and *cgh-1* gene products by facilitating the decay of their mRNAs. Published bulk and single-cell RNA-seq datasets on *C. elegans embryos* suggested that *ifet-1, car-1,* and *cgh-1* transcripts are maternally deposited and subjected to rapid degradation before MZT^77,78^. Consistent with our hypothesis, an initial RT-qPCR on mixed-stage embryos revealed a 1.4 and 1.6-fold increase in *car-1* and *cgh-1* mRNAs, respectively, in *edc-3(0)* in comparison to WT, and this increase was comparable (1.6 and 2-fold) in the *edc-3(0); edc-4(0)* double mutant **(Figure S5A)**. Meanwhile, *edc-4(0)* expressed WT levels of *car-1* and *cgh-1* mRNAs, in agreement with EDC-4 being dispensable for their somatic clearance **(Figures 3, S3)**.

To determine the timeline and lineage (somatic or germ cells) wherein EDC-3 promotes embryonic mRNA degradation, we performed single-molecule fluorescence in situ hybridization (smFISH)^79,80^ using probes against the *cgh-1* mRNA coding sequence to trace and quantify *cgh-1* mRNAs in early embryos. smFISH was carried out in WT, *edc-3(0)*, *edc-4(0),* and *edc-3(0); edc-4(0)* double mutant embryos co-expressing the germ cell marker PGL-1::GFP **(Figures 6A, S5)**. In WT embryos, total *cgh-1* mRNA counts increased by 7% (on average 891 more mRNA molecules per embryo) between 2 and 9-cell embryos **(Figure 6B)**. This increase was primarily driven by the somatic pool of *cgh-1* mRNAs, which was increased by 22% (on average 2,558 mRNA molecules), presumably a result of zygotic transcription of *cgh-1* following zygotic activation at ∼4-cell **(Figure 6C)**. The somatic pool of *cgh-1* mRNAs per embryo underwent a 92% reduction (a reduction of 10,850 RNA molecules) from 9 to >100-cell embryos, consistently with the degradation of maternal and possibly also zygotic *cgh-1* mRNAs **(Figure 6C)**. The distribution of mRNAs also changed with development; *cgh-1* mRNAs were concentrated in 50-70 clusters in the soma of 2 to 16-cell embryos, but nearly 100% of these clusters were resolved in 33∼66-cell embryos **(Figure 6D)**. In the WT germline, *cgh-1* mRNAs remained enriched in and largely associated with germ granules (PGL-1-positive foci) from 2 to ∼100-cell (after the P4 blastomere divided into the Z2 and Z3 germline precursors), and diminished shortly after morphogenesis **(Figure 6A upper panel)**. The slower decay of *cgh-1* mRNAs in germline lineage in comparison to somatic blastomeres agrees with the reported delayed onset of MZT in the germline, and the proposed role of germ granules to protect maternal mRNAs from degradation^54,81,82^.

**Figure 6.**
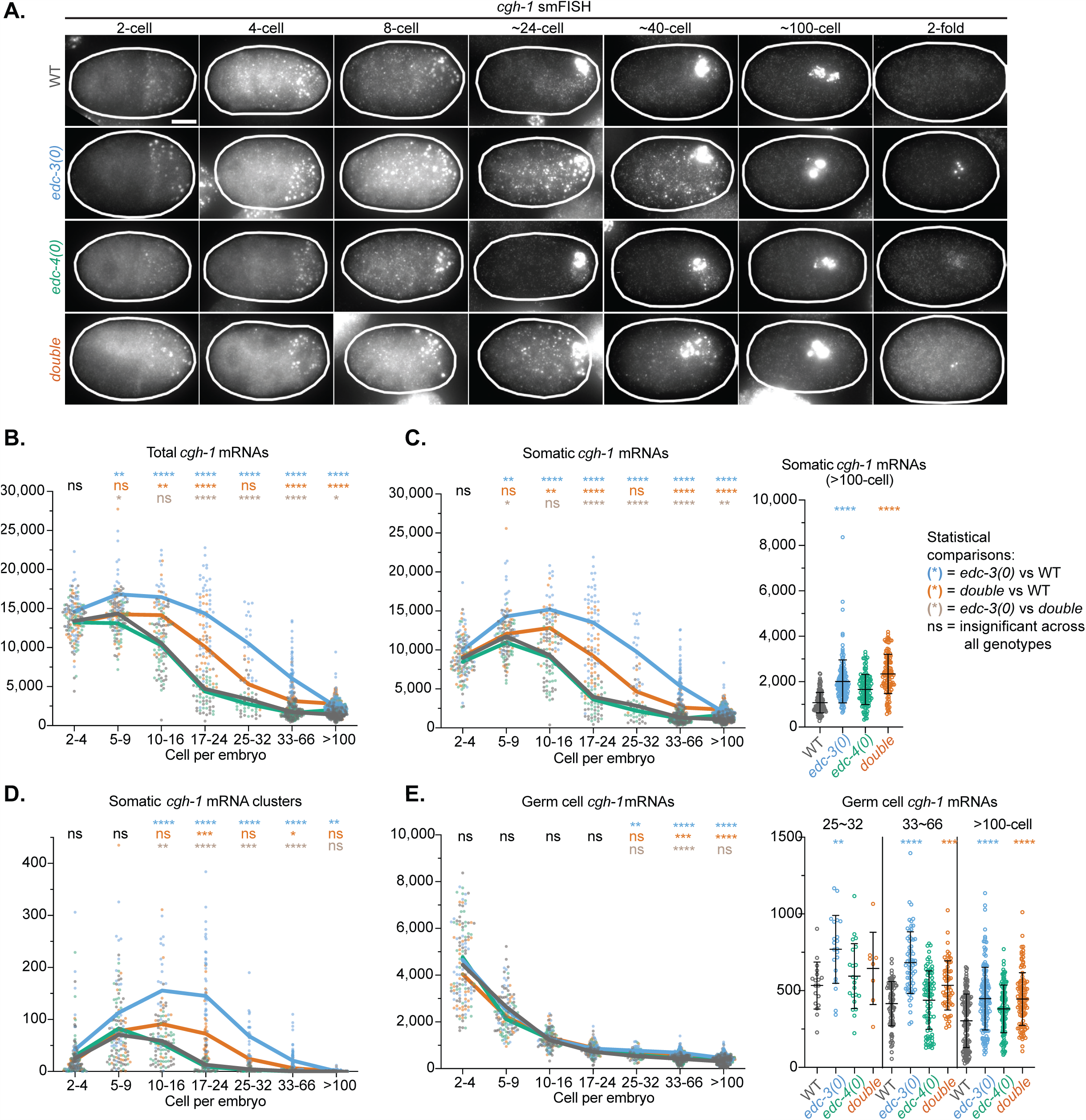
EDC-3 promotes the somatic and germline clearance of *cgh-1* mRNAs. (A) Representative photomicrographs from single-molecule Fluorescence *In Situ* Hybridization (smFISH) to detect *cgh-1* mRNAs on WT, *edc-3(0)*, *edc-4(0)*, and *edc-3(0); edc-4(0)* double mutant embryos. Scale bar is 10 µm. The corresponding DAPI and PGL-1::GFP (germ cell marker) images are shown in **Figure S5C**. (B) Total *cgh-1* mRNA counts per embryo over embryonic development. (C) Somatic *cgh-1* mRNA counts per embryo over development. (Right) Zoom of >100-cell embryos to better visualize differences. (D) Somatic *cgh-1* mRNA cluster counts per embryo over development. (E) Germ cell *cgh-1* mRNA counts per embryo over development. (Right) Zoom of data from >25-cell embryos to better visualize differences. Results are representative of three technical and biological replicates. Between 2-60 embryos from each stage bin from each genotype were analyzed per replicate (on average 15 embryos per stage bin). Data points are coloured as follows: WT in dark grey, *edc-3(0)* in blue, *edc-4(0)* in green, *edc-3(0); edc-4(0)* in orange. In each stage bin, Tukey’s multiple-comparisons test was conducted. Statistical analyses were denoted in colour as follows: Blue = *edc-3(0)* vs WT; Orange = *edc-3(0); edc-4(0)* double mutant vs WT; Tan = *edc-3(0); edc-4(0)* double mutant vs *edc-3(0)* single mutant; ns in black indicates insignificant difference across all genotypes. p-values: (*) =0.01-0.05, (**) =0.001-0.01, (***) =0.0001-0.001, (****) =<0.0001, p≥0.05 are insignificant (ns). **See also Figure S5.**

In comparison to WT, *edc-3(0)* embryos progressively accumulated up to 3.5-fold more *cgh-1* mRNA molecules in total (when considering the soma and germline altogether) from 5 to 66-cell, and 1.8-fold more than WT in embryo cohorts of >100-cell, supporting the promotion of *cgh-1* mRNA degradation by EDC-3 **(Figure 6B)**. Notably, in embryo cohorts of 5 to 16-cell, wherein *cgh-1* mRNA started to decline in WT, somatic *cgh-1* mRNA counts still increased by 6% (881 mRNA molecules) in *edc-3(0)* embryos (**Figure 6C)**. The difference with WT was further accentuated in *edc-3(0)* embryo cohorts of 17 to 66-cell, wherein ∼3-fold more total mRNA molecules were counted than in WT. *cgh-1* mRNAs were also more concentrated in the soma of *edc-3(0)* embryos; up to 20 times more clusters were detected than in WT among the 17 to 66-cell embryo populations **(Figure 6D)**. While overall *cgh-1* mRNA stabilization in *edc-3(0)* was primarily driven by impaired degradation of the somatic pool in all embryonic stages, a significant 1.4 to 1.6-fold accumulation of *cgh-1* mRNAs in *edc-3(0)* germ cells was detected in embryo cohorts of 25-cell and beyond **(Figure 6E, right panel)**. It is noteworthy that this coincides with the onset of maternal mRNA decay in the germline^54^. Furthermore, more concentrated *cgh-1* mRNA clusters were observed in Z2 and Z3 of *edc-3(0)* in comparison to WT, and on average 1.7 times more germline *cgh-1* mRNA clusters persisted in *edc-3(0)* than WT at >100-cell, which included the morphogenetic stages **(Figure 6A, ∼100-cell and 2-fold; Figure S5B)**. Lastly, we did not observe any difference in PGL-1 perinuclear distribution in *edc-3(0)* **(Figure S5C)**.

In contrast *to edc-3(0)*, *cgh-1* mRNA counts in *edc-4(0)* mutant embryos did not differ from WT across all embryonic stages in both soma and germline blastomeres. Interestingly, in double *edc-3(0); edc-4(0)* mutant embryos, *cgh-1* mRNA degradation was also delayed, and mRNAs were clustered, but this stabilization was partially suppressed in both soma and germ blastomeres across all stages when compared to *edc-3(0)* alone **(Figure 6, S5B-E)**.

These data indicate that EDC-3 promotes *cgh-1* mRNA decay in both somatic and germ cell blastomeres and regulates its concentration into clusters. We note, however, that we cannot infer whether the clustering of *cgh-1* mRNAs into foci is a cause or consequence of their delayed degradation in *edc-3(0)*. Overall, our data implicate the decapping scaffold EDC-3 in the decay of mRNAs through embryonic development from the onset of zygotic transcription to morphogenesis, and support an antagonistic role for EDC-4 in these functions (See Discussion).

## Discussion

This work delineates the roles and interactions of the DCAP-2 scaffold proteins EDC-3 and EDC-4 in the embryonic development of *C. elegans*. Through quantitative imaging, we find that EDC-3 facilitates the removal of embryonic *cgh-1, car-1*, and *ifet-1* mRNAs. This not only reduces their expression but also counters the formation of DCAP-2 biomolecular assemblies in the developing somatic cells and, subsequently, in the germline. EDC-4 counteracts EDC-3 in these functions and further recruits the GID complex, a ubiquitin ligase known for its involvement in MZT in other species, to DCAP-we2. Lastly, we uncover a novel role for EDC-3 in defining the boundaries among distinct biomolecular condensates, specifically P-bodies, germ granules, and stress granules.

### EDC-3 promotes the timely clearance of embryonic mRNAs

Our findings reveal a novel role for EDC-3 in the clearance of maternal transcripts within both somatic and germline domains, which operates within a unique developmental timeframe that complements other embryonic mRNA turnover mechanisms **(Figure 7)**. In *C. elegans*, maternal mRNA clearance initially occurs during the oocyte-to-embryo transition, facilitated by polyC motifs in certain mRNA 3’-UTRs and endogenous siRNA activities^83^ **(Figure 7A)**. A subsequent wave of clearance, involving both maternal and zygotic proteins, begins with zygotic transcription around the 4-cell stage^54,84^. This phase primarily involves the Argonaute CSR-1, which, guided by maternally inherited 22G-RNAs, targets about half of the transcripts for degradation in its regulatory domain **(Figure 7B)**^85^. However, this activity is predominantly observed in embryos from the 4-cell to the 20-cell, after which CSR-1 expression in the soma diminishes^85^. Additionally, members of the AU-rich element binding, CCCH finger family of proteins, MEX-3, MEX-5 and MEX-6, promote the decay of specific maternal transcripts (*nos-2*, *cpg-2*, *chs-1*) in the soma, also up to ∼20-cell, after which the MEX proteins are depleted from the soma and remain enriched in the germline lineage^4,86,87^.

**Figure 7.**
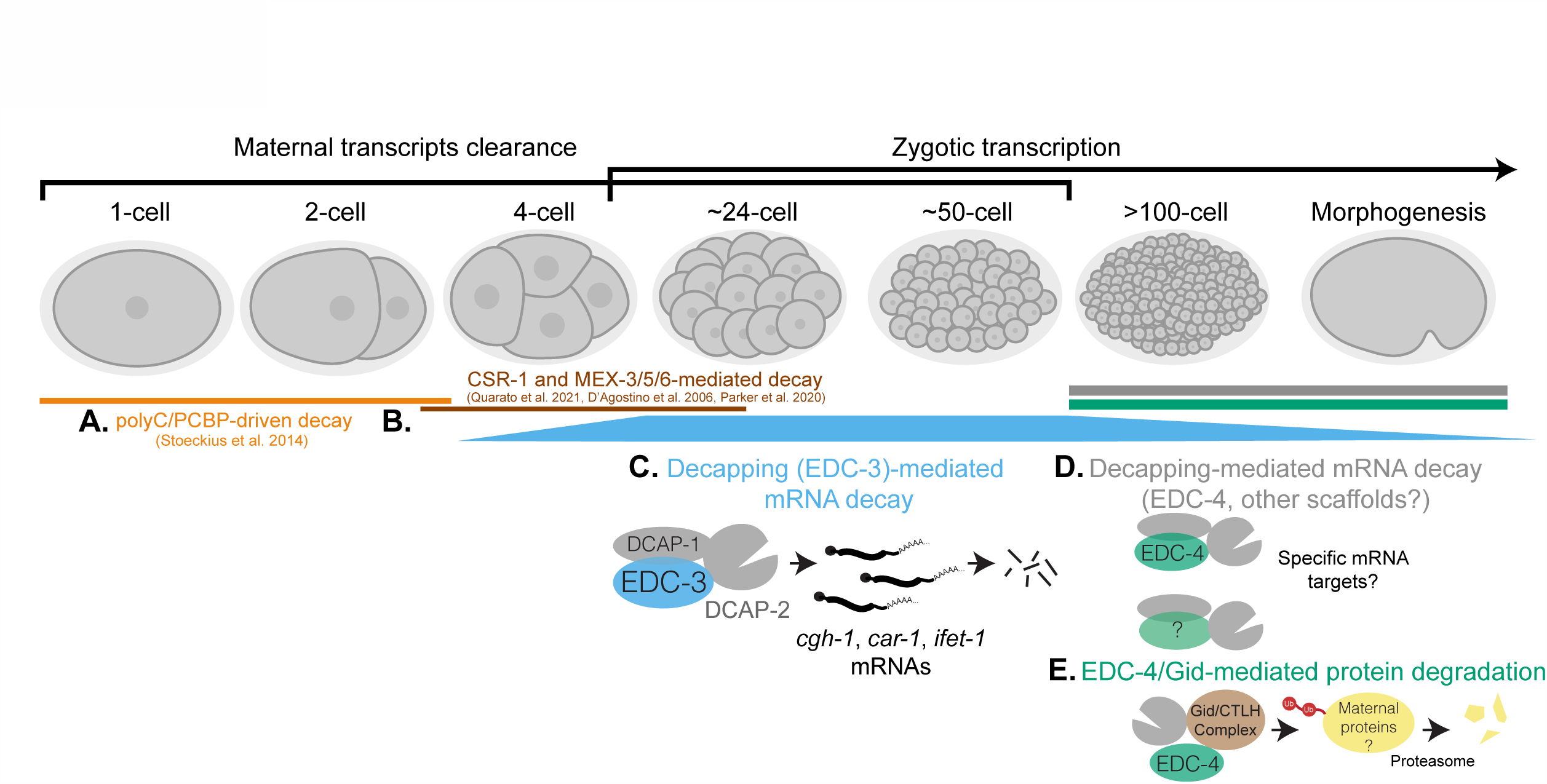
Model of EDC-3 and EDC-4 functions in the decay of embryonic mRNAs and proteins. (A) During oocyte-to-embryo transition, the bulk of maternal mRNAs are cleared in a polyC-driven mechanism. (B) CSR-1 and some members of the MEX family proteins promote the decay of maternal transcripts, after which CSR-1 and MEX expression decline dramatically in the soma. (C) EDC-3 mediated decapping promotes the somatic and germline clearance of *cgh-1*, *car-1*, *ifet-1*, and possibly other maternal and zygotic mRNAs, at post-4-cell embryos up to early morphogenetic stages. (D) EDC-4 and other unidentified decapping scaffolds promote the decapping and decay of embryonic mRNAs after the decline of EDC-3’s role and expression in late embryonic stages. (E) Recruitment of the Gid ubiquitin ligase complex to DCAP-2 by EDC-4 may regulate the degradation of maternal proteins.

Our data demonstrates that EDC-3 plays a pivotal role in the decay of maternal transcripts around the domain of, and following the reduction in somatic CSR-1 and the MEX family proteins **(Figure 7C)**. This is supported by our smFISH data showing the removal of *cgh-1* mRNAs between the 4 and ∼100-cell stages in WT embryos. Conversely, in *edc-3(0)* embryos starting from >4-cell and most predominantly between 8 and ∼100-cell, *cgh-1* mRNAs, as well as DCAP-2 and IFET-1 foci, persist abnormally **(Figure 6)**. When considered with our detailed profiling of DCAP-2 expression, these results suggest a dynamic engagement of DCAP-2 in embryos. DCAP-2 expression is minimal in early embryos and increases significantly post-4-cell **(Figure 1A)**, peaking as somatic CSR-1 and MEX decline after ∼20-cell, indicating that the influence of decapping and decay in the soma rises with zygotic contribution and persists through the cessation of CSR-1 and MEX activities.

Our findings further support a role for EDC-3 in mRNA decay in the germline, as evidenced by the persistence of *cgh-1* mRNAs in the Z2 and Z3 primordial germ cells of *edc-3(0)* mutants **(Figure 6A)**. Low levels of DCAP-2 in the germline were observed in this study **(Figures 3, S3)** and by others^88^. Meanwhile, IFET-1 is enriched in and associated with germline P-bodies up to ∼50-cell stage, beyond which it becomes undetectable **(Figures 3, S3)**. The distinct expression patterns and ratios of these factors suggest that EDC-3’s germline mRNA decay role might involve a pathway different from the ones that prevail in somatic cells. This in turn raises the possibility that CSR-1 and the MEXs, along with other yet-to-be-identified factors, may utilize decapping co-activators as scaffolds for the selective degradation of maternal transcripts in the germline. Despite the potential for mechanistic parallels with the soma, mRNA decay during maternal-to-zygotic transition (MZT) in the germline remains significantly less understood than in somatic cells. It will thus be interesting to identify which trans-acting factors harness EDC-3 not only in the soma but also in the germline.

### Decapping scaffolds are functionally specialized in *C. elegans* development

Prior evidence would have led to the expectation that decapping scaffold proteins EDC-3 and EDC-4 function redundantly or compensate for each other in controlling DCAP-2 activity and biomolecular assemblies. Our results instead support the view that decapping scaffolds are specialized, distinctly regulated, and can even antagonize each other. EDC-3 and EDC-4 have very different domains of expression along embryonic and larval development; whereas EDC-3 is highly enriched in embryos and decreases dramatically at L2, EDC-4 expression is nearly steady across development **(Figure 4F)**. Functionally, we show that EDC-3 and EDC-4 have opposing functions on assemblies of DCAP-2 in the embryo; EDC-4 is required for the formation of large foci while EDC-3 limits these assemblies **(Figure 1)**, and conversely, while *edc-3(0)* leads to persistence of *cgh-1* mRNA, *edc-4(0)* partially rescues its timely clearance in *edc-3(0)* **(Figure 6)**.

Further along the lines of their functional specialization, EDC-3 and EDC-4 only partially share their engagement with embryonic miRNAs **(Figures 4C-E)**. Whereas *dcap-*loss strongly interacts genetically with embryonic and larval miRNA hypomorphs, *edc-*seems to selectively enable *lsy-6* function in left-right asymmetry, which occurs during morphogenesis^71^. In contrast, *edc-4* does not partake in this embryonic miRNA’s function, and both *edc-3 and edc-4* exacerbate a *miR-35-42* family hypomorph. Interestingly, neither *edc-3* nor *edc-4* is required for *let-7* function during the L4-to-adult transition, despite a clear genetic interaction between *let-7* and *dcap-2*. This raises the possibility of a role for yet another decapping protein scaffold during larval development **(Figure 7D)**. As of now, its identity is unknown, but we ruled out the PATR-1 protein as a candidate, as its depletion did not significantly affect the *let-7* hypomorph (not shown).

The distinct functions of EDC-3 and EDC-4 in embryogenesis are also reflected through their importance for the molecular commitments of DCAP-2 in its interactome **(Figure 2)**. For example, EDC-4 promotes DCAP-2 interaction with the GID complex **(Figure S2D)**, thus providing a novel physical linkage between the control of mRNA and protein degradation during MZT **(Figure 7E)**. Our data further indicate that the GID complex is dispensable for the degradation of IFET-1 **(Figure 5)**, in stark contrast with *Drosophila* embryo^73,74^. We reason that this interaction may target other maternal proteins in *C. elegans,* likely later in development, in line with the expression domain of the complex **(Figure 7E)**. This mechanism could complement other maternal protein degradation pathways, such as the targeting of maternal meiotic and germ plasm proteins by the Cullin ubiquitin ligase^89,90^ and autophagy^91,92^.

### EDC-3 enforces boundaries of biomolecular condensates

Another key finding of this study is that the loss of EDC-3 triggers a partial convergence of P-bodies, stress granules, and germ granule interactomes on DCAP-2, highlighting EDC-3’s role in managing the composition of biomolecular condensates **(Figures 2B, S2C)**. We propose that EDC-3 might mediate this by modulating the interaction between the IFET-1/CAR-1/CGH-1 complex and DCAP-2 in P-bodies, rather than directly sorting or restricting proteins within condensates. This hypothesis is supported by several lines of evidence. First, CGH-1 and its human ortholog Ddx6, interact with stress and germ granule proteins^93,94^. As CGH-1 is upregulated upon loss of *edc-3*, this may thus explain their enhanced interactome footprint with DCAP-2 **(Figure 3G)**. Additionally, CGH-1 has been identified as a key factor in preventing the merging of P-bodies and stress granules by acting as a bridge protein, potentially through its activity in RNA helicase-mediated mRNP remodelling^20,34,93,95^. One possibility is that EDC-3 may indirectly influence biomolecular condensate interactions via CGH-1’s enzymatic function. Unfortunately, as loss of *cgh-1* is lethal^45,46^, we could not directly assess its impact on the interactomes of EDC-3 or DCAP-2. Furthermore, the reduction in DCAP-2 foci in *edc-3(0)*; *edc-4(0)* mutants upon RNAi against components of the IFET-1/CAR-1/CGH-1 complex **(Figures 2D-F)**, indicates that this complex is crucial for forming the molecular framework of embryonic condensates. As this genotype *(edc-3(0); edc-4(0))* also partially mitigates the delay in mRNA clearance observed in *edc-3(0)* **(Figure 6)**, the ratio of IFET-1/CAR-1/CGH-1 complex relative to EDC-4 may influence decapping and decay activities, regardless of their occurrence within or outside of biomolecular condensates.

### Limitations of the study

This work, like virtually all in the mRNA decay or biomolecular condensate fields, did not directly establish whether mRNA decapping and decay occur inside, or physically associate with P-bodies. The fact that *edc-3(0)* leads to an increase in P-body size, while at the same time delays mRNA clearance, could be interpreted as support against such a straightforward association. Perhaps a key nuance brought to light by this study is that even if they share a ‘core’ or ‘signature’ marker including DCAP-2 itself, biomolecular condensates composition can differ widely with cellular identity or perturbations to interaction networks. Consequently, the activities of P-bodies may vary depending on their composition. Another challenge is defining the full range of mRNA targets regulated by EDC-3. We reason that the regulome of EDC-3 likely includes a subset of specific maternal transcripts, the identity of which may be revealed by the identification of trans-acting factors and their associated elements. Advancements in single-cell sequencing technology could enable quantitative spatiotemporal profiling of EDC-3 targets featured in this work. Lastly, P-bodies and germ granules have been involved in other gene silencing mechanisms such as translation repression and transgenerational epigenetic inheritance^94^. These processes likely operate concurrently or intersect within gene regulatory networks. Future efforts aimed at clarifying how these mechanisms are coordinated, or function independently are likely to be insightful.

## Acknowledgments

This work was supported by the Canadian Institute of Health Research (CIHR) grants (MOP-123352) to T.F.D. We thank Dr. Marc Fabian, Dr. Martin Simard, and Hsin-Wei Tseng for constructive feedback on the manuscript. Live imaging data was acquired in the McGill University Advanced BioImaging Facility (ABIF), RRID:SCR_017697 with training and analysis support from Dr. Joel Ryan. We thank Dr. Abigail Gerhold for training E.V. on live imaging and for providing ImageJ macros for image registration. The following strains were provided by the Caenorhabditis Genetics Center (CGC), which is funded by the NIH Office of Research Infrastructure Programs (P40 OD01044): *dcap-2(ok2023)*, *pgl-1(gg547)*, *ntIs1*, *lsy-6(ot150)*, *let-7(n2853)*, *pgl-1(ax3122)*. We thank Dr. Katherine McJunkin for providing the *mir-35-41(nDf50)* strain, Alice Lambert for providing the plasmid for CGH-1 purification and antibody production, and Douna Husin for assisting with the generation of *dcap-2(qe79)* strain. E.V. was supported by the Canderel Entrance Studentship, J.P. Collip Studentship, Charlotte & Leo Karassik Foundation Studentship, and the Donner Foundation Studentship.

## Author contributions

E.V. and T.F.D. conceived the study and designed experiments. E.V. performed all experiments and analyzed data with the following exceptions: Y.J.A. and A.M. performed mass spectrometry experiments, J.R. and J.M.S.X. provided codes for smFISH analysis, T.C. performed *miR-35-42* analysis, R.L. helped generate some strains. T.F.D., M.V., J.W., N.S. supervised and provided input on experimental design and analyses. E.V. prepared figures with input from all authors. E.V. and T.F.D. wrote the manuscript with input from all authors.

## Methods

### Lead Contact and Materials Availability

Further information and requests for resources and reagents should be directed to and will be fulfilled by the Lead Contact, Thomas F. Duchaine

### Experimental Model and Subject Details

1. *C. elegans* were maintained using standard procedures at 16°C unless noted otherwise and fed OP50 *E. coli* ^96,97^. The strains used in this study are listed in the **Table S2** (Strains tab).

### Method Details

#### CRISPR/Cas9

Gene editing was done using Cas9 ribonucleoprotein and *dpy-10* co-conversion strategies as previously described^49,98^. sgRNAs (fused crRNA and tracrRNA) were generated by extending a common sgRNA(F+E) sequence with gene-specific crRNA sequence downstream of a T7 promoter by PCR with Phusion polymerase (NEB). PCR product was gel extracted and sgRNA was *in vitro* transcribed using the MegaShortScript™ T7 Transcription Kit (Invitrogen AM1354) at 30°C overnight, DNase-treated, extracted with phenol chloroform, precipitated with isopropanol, and dissolved in water. For addition of small epitope tags (FLAG/Myc/V5/HA) or introduction of point mutations, ssODN repair templates were synthesized by IDT. To generate *dcap-2::gfp(S65C, Q80R)* and *ifet-1::TagRFP,* repair templates were generated by amplifying genomic DNA from OH3192 (*ntIs1*(*gcy-5p::GFP* + *lin-15(+) V*)) and YY968 *(znfx-1(gg544(3xflag::gfp::znfx-1); pgl-1(gg547(pgl-1::3xflag::tagRFP*), respectively, using primers complementary to GFP/TagRFP with added overhangs to introduce homology arms. Multiple PCR reactions were pooled and concentrated using QIAQuick PCR purification kit (Qiagen 28104) to reach a minimum of 1µg/µL concentration. Injection mix contains 150mM KCl, 10mM HEPES pH 7.4, 0.183µg/µL *dpy-10* sgRNA, 16.75ng/µL dpy-10 ssODN repair template, 533ng/µL gene-specific sgRNA, 146.7ng/µL gene-specific ssODN repair template (for small edits) or 292ng/µL gene-specific PCR repair template (for large edits), and 2.4 µg/µL in-house purified Cas9 protein^49^. The mix was injected into both germlines of *C. elegans* fed with HT115 bacteria expressing *cKu80* RNAi^48^ using standard protocol^99^. Dumpy and Roller F1s were screened by PCR and homozygotes for the edit were isolated and sequenced. All CRISPR-generated strains were outcrossed twice. Oligonucleotides to produce sgRNA and repair templates are listed in **Table S2** (Oligonucleotides tab).

#### Antibody production

Sequences encoding full-length EDC-3, CGH-1 and a ∼37 kDa fragments of EDC-4 (sequence: NAEDSGDDLF-(314 amino acids)-QKCLVSLVSQ) were individually cloned into pSMT3 plasmid (N-terminal 6XHis-SUMO). Each construct was expressed in and purified from BL21(DE3) cells. For expression, transformed colony was grown to OD_600_ of 0.4-0.6, induced overnight with 1 mM IPTG, and pelleted for purification. Cells were lysed and sonicated in 6 M guanidium hydrochloride, 20 mM sodium phosphate, 0.5 M NaCl, and protease inhibitor (Sigma P8340) at pH 7.4. The soluble fraction was bound to Ni Sepharose 6 Fast Flow resin (Cytiva 17-5318-01), washed twice with wash buffer (8 M urea, 20 mM sodium phosphate, 0. 5 M NaCl, 60 mM imidazole) and eluted with elution buffer (8 M urea, 20 mM sodium phosphate, 0.5 M NaCl, 250 mM imidazole). Protein-containing fractions were pooled and concentrated with Amicon Ultra-15 centrifugation filters. Two mg of proteins were used in four rounds of rabbit injection (Capralogics, Cambridge Massachusetts) and a total of three bleeds were collected and characterized.

#### Immunoprecipitation (IP)

Embryos were collected using standard bleaching procedure from 15-cm NMG plates containing ∼50,000 gravid adults per plate grown from synchronized/hatched L1s, and flash-frozen in liquid nitrogen^97^. Frozen embryo pellet was resuspended in 2-3X volumes of lysis buffer (50 mM Tris-Cl pH 7.4, 150 mM NaCl, 1 mM EDTA pH 8.0) with cOmplete™ protease inhibitor (Roche 4693159001) and Phosstop™ phosphatase inhibitor (Roche 4906845001). The mixture was homogenized 40x using a Dounce homogenizer on ice, clarified 3x by 17,000 g centrifugation for 10 minutes and quantified. For LC-MS/MS, 5 mg of proteins at 2.5 mg/mL was used. For Co-IP and western blotting, 2-2.5 mg of proteins was used. Lysates were treated with 100 mg/mL RNase A (Thermo Scientific EN0531) for 1 hour at 4°C and centrifuged at 17,000 g for 2 minutes, and the supernatant was subjected to IP with 10 µL of packed anti-FLAG M2 magnetic beads (Sigma-Aldrich M8823) per mg proteins for 2 hours at 4°C. For Figure 4F specifically, anti-HA (C29F4) beads (Cell Signaling Technology 3724S) were used. The beads were washed 3X with lysis buffer. For Co-IP and western blotting, proteins were eluted from beads with SDS loading buffer and heated at 95°C for 2 minutes. For LC-MS/MS, beads were further washed 3X with TBS (150 mM Tris-Cl pH 7.4, 150 mM NaCl) and eluted 3X with 100 µL of freshly prepared 0.5 M ammonium hydroxide. Pooled eluates were concentrated by speed-vac centrifugation, washed with nuclease-free water, and finally dried by speed-vac before analysis by LC-MS/MS.

#### Protein Identification by LC-MS/MS

Dried IP eluates were resuspended in 100 mM Tris-HCl pH 8.5, 8 M urea, then reduced and alkylated using 5 mM Tris (2-carboxyethyl) phosphine and 10 mM iodoacetamide, respectively. The samples were proteolytically digested with Lys-C and trypsin overnight at 37°C. The digestion was quenched with 5% formic acid and desalted either using C18 tips (Thermo Fisher 87782) or equal mix of Sera-Mag Carboxylate SpeedBeads (Cytiva 65152105050250 & 45152105050250)^100^. Dried peptides were resuspended in 5% formic acid and analyzed by LC-MS/MS. Briefly, peptides were separated by reversed-phase chromatography using 75 µm inner diameter fritted fused silica capillary column packed in-house to a length of 25 cm with Luna C18 3µm reverse phase particles^101^. The increasing gradient of acetonitrile was delivered by a Dionex Ultimate 3000 (Thermo Scientific) at a flow rate of 300 nL/min. MS/MS spectra were collected using data-dependent acquisition on Orbitrap Fusion Lumos Tribrid mass spectrometer (Thermo Fisher Scientific) with an MS1 resolution (r) of 120,000 followed by sequential MS2 scans at a resolution (r) of 15,000. The data generated by LC-MS/MS were analyzed on MaxQuant bioinformatic pipeline^102^. The Andromeda component of MaxQuant was employed as the peptide search engine and the data were searched against *Caenorhabditis elegans* reference proteome (Uniprot Reference UP000001940). A maximum of two missed cleavages was allowed and the maximum false discovery rate for peptide and protein was specified as 0.01. Label-free quantification (LFQ) was enabled with LFQ minimum ratio count of 1. The parent and peptide ion search tolerances were set as 20 and 4.5 ppm respectively. The MaxQuant output files were processed for statistical analysis of differentially enriched proteins using ProVision^103^, which adapted the Limma package^104^. Analysis was done on the log2 transformed LFQ intensities of proteins with at least 2 unique peptides. Three biological replicates from each genotype were analyzed against the corresponding N2 (untagged). Proteins found in at least 2 biological replicates in at least 1 group were kept for analyses. Missing values were imputed with default parameters (width = 0.3, downshift = 1.8). Proteins with p-value (Benjamini Hochberg FDR-corrected) < 0.05 and log2FC > 1.8 were deemed significant interactors.

#### Western blotting

Antibodies generated in this study and used in western blots were: anti-EDC-4 (Rabbit #6753 production bleed 1:1000), anti-EDC-3 (Rabbit #7503 production bleed 1:5000), anti CGH-1 (Rabbit #7199 production bleed 1:2000) (Capralogics). Commercial primary antibodies used were listed in **Table S2** (Antibodies tab).

Secondary antibodies used were: IRDye 800CW Goat anti-Rabbit (1:10 000) (LI-COR), IRDye 680RD Goat anti-Mouse (1: 10 000) (LI-COR). Fluorescent TrueBlot® Rat anti-Mouse IgDyLight™ 680 (1: 1000) (Rockland) was used when expected bands were of similar size to IgG in Co-IP experiments.

Proteins were transferred from SDS PAGE onto Immobilon-FL PVDF membranes (Millipore Sigma) using a semi-dry transfer apparatus (15V, 0.2A, 1 hour), blocked for 1 hour in Odyssey or Intercept blocking buffer (LI-COR), then incubated with primary antibody dilution overnight at 4°C. Membranes were washed three times with PBST, incubated with secondary antibody dilutions at room temperature for 1 hour, washed three times with PBST, and scanned on a LI-COR scanner.

#### RT-qPCR

Total RNA was extracted from ∼20 µL of frozen embryos using 1mL TRIzol™ (Invitrogen 15596026) and subsequently treated with TURBO™ DNase (Invitrogen AM2238) for 1 hour at 37°C. Reverse transcription (RT) was performed on 1 µg RNA with the iScript™ Reverse Transcription Supermix (Bio-Rad 1708841) for 5 min at 25°C, 30 min at 42°C and 1 min at 95°C. cDNA was diluted four-fold and 2 µL was used for quantitative PCR (qPCR) using SsoAdvanced Universal SYBR Green Supermix (Bio-Rad 1725274) and 500 nM of gene-specific qPCR primers (see **Table S2**, Oligonucleotides tab). qPCR was done on a Bio-Rad CFX Connect™ Real-Time PCR Detection System with the following parameters: initial denaturation at 95°C for 2 min and 40 cycles of 15 s at 95°C, 59°C for 30 s and 72°C for 30, and final melt curve of 65C to 95C in 0.5C steps at 5 s per step. Fold change gene expression was calculated using ΔΔCt method, normalized to *act-1*.

#### RNAi

*cgh-1* and *ifet-1* RNAi were performed by feeding L4 animals with HT115 expressing double-stranded RNA^105^. The gene sequence was PCR-amplified with primers to introduce T7 promoters on both ends (see **Table S2**, Oligonucleotides tab) and cloned into pSC-A-amp/kan (Stratagene) vector. Sequences containing an 80 bp portion of L4440 plasmid’s multiple cloning site flanked by T7 promoters were used as a negative control, i.e. *ctrl(RNAi)*. Plasmids containing genes to be knocked down or negative control sequences were transformed into HT115 and plated on ampicillin and tetracycline-containing media. A single colony was cultured overnight in 1 litre LB containing 50 μg/mL ampicillin, pelleted and diluted in 5X volumes of M9. The diluted RNAi bacteria was seeded on NGM plates containing 50 μg/mL ampicillin and 1 mM IPTG, and double-stranded RNA expression was induced at 37°C for 4.5 hours. For analyses, one L4 animal was picked onto each plate and the F1s were analyzed. Progenies from at least 5 plates were analyzed in each experimental replicate.

#### Live Imaging

Embryos were dissected in M9 buffer from gravid adults, transferred to a 2% agarose pad on a glass slide and covered with a #1.5 coverslip. Coverslip edges were sealed with VaLaP (1:1:1 Vaseline, lanolin, paraffin), backfilled with M9 buffer, and imaged at room temperature (∼20°C) immediately, within <10 minutes between embryo mounting and the end of image acquisition, to minimize potential imaging stress-induced effects on condensates^106^. All live images were acquired using a 63X 1.40 NA objective on a Leica DMI60000B inverted microscope equipped with a Quorum WaveFX-X1 spinning disk confocal system, with an ASI MS-2000 piezo stage, and two Hamamatsu “ImagEM” EM-CCD cameras, controlled by the MetaMorph software. For DCAP-2::GFP, acquisition used 200 ms exposure time with 491 nm (50 mW) diode laser at 25% intensity and ET 525/50 emission filter. For IFET-1::TagRFP, acquisition used 200 ms exposure time with 568 nm (50mW) diode laser at 5% intensity and ET 620/60 emission filter. Gain was set at 2 and no binning was applied. For GFP/RFP co-imaging, samples were excited simultaneously using dual camera setting following confirmation of no spectral bleed-through by imaging each GFP/RFP-only strain using the same settings. Images of 1 µm TetraSpeck microspheres were acquired in parallel for image registration (0.1 µm step size, 100 ms exposure time, 7% and 15% laser intensity on GFP and RFP, respectively). For Figure 1, 25 confocal sections with a 1.0 µm step size over 24 µm depth of the embryos were acquired. For subsequent analyses, 36 confocal sections with a 0.7 µm step size were acquired. One slice of brightfield image was acquired from the middle plane of the embryos for staging.

#### Live Imaging Analysis, Foci Detection and Quantification

Embryos were outlined manually using the polygon selection tool on Fiji^53^ and staged using brightfield images as a reference. A median filter of 1 pixel was applied to each confocal stack to reduce noise. Total intensity per embryo was measured by taking the mean intensity of sum slice projection from all stacks. Background intensity was acquired from non-fluorescent embryos imaged with the same acquisition parameters and subtracted from the raw mean intensity. To detect DCAP-2 and IFET-1 foci, the maximum intensity projection from all stacks was acquired. Automatic foci segmentation was done using the StarDist 2D plugin on Fiji^51,52^ using a probability threshold of 0.2 and overlap threshold of 0.4 (for DCAP-2) or 0.2 (for IFET-1). The regions of interest (ROIs) of the foci were applied to the median-filtered maximum projection images and the mean intensity and Feret’s diameter of each spot were measured using the analyze particle function. Background intensity for foci was acquired from an empty area outside of the embryo and subtracted from the raw mean intensity. The averages of mean intensity and diameter of all foci in an embryo were plotted. For all intensity measurements, differences across genotypes were represented as fold change relative to WT at 1∼2-cell or 4∼6-cell. Representative images were maximum projections of median-filtered images.

#### Single Molecule Fluorescence In Situ Hybridization (smFISH)

smFISH was performed as previously described^80^. Briefly, embryos were harvested from 2x15-cm dishes of synchronized gravid adults by standard bleaching procedures, fixed with 1 mL cold methanol, freeze-cracked for 1 minute in liquid nitrogen, and repeatedly centrifuged for up to 4 minutes. Methanol was replaced with cold acetone, incubated for 3 minutes, and repeatedly pelleted for up to 2 minutes. Fixed embryos were washed in 1 mL Stellaris Wash Buffer A (LGC Biosearch SMF-WA1-60) with 10% formamide (Sigma F9037) for 5 minutes and hybridized with 100 μL of Stellaris Hybridization Buffer (LGC Biosearch SMF-HB1-10) containing 22.3 nM smFISH probes (48 probes targeting *cgh-1* coding sequence conjugated with Quasar 670 dye, see **Table S2**, smFISH Probes tab) for ∼16 hours in the dark with shaking at 37°C. Hybridized embryos were washed with 1 mL Wash Buffer A with 10% formamide for 30 minutes at 37°C, counterstained with 1 mL Wash Buffer A supplied with 1 ng/μL DAPI for 30 minutes at 37°C. Pelleted embryos were washed with 1 mL Stellaris Wash Buffer B (LGC Biosearch SMF-WB1-20) for 5 minutes at RT and incubated in 20-30 μL mounting medium (50% glycerol, 20 mg/mL n-propyl gallate, 80 mM Tris pH 8.0) at 4°C for 30 minutes. Two μL of embryo mixture was mixed with 2 μL of VECTASHIELD (Vector Labs H-1000-10) on an 8-mm round cover glass #1.5, sandwiched with 22x22-mm square cover glass #1.5, and fixed onto a microscope slide using the Press-To-Seal Silicone Isolator (Grace Bio-Labs JTR20R-0.5). Slides were imaged on a Nikon Eclipse Ti-2 widefield microscope with a Spectra X LED light engine (Lumencor), and an Orca-Fusion sCMOS camera (Hamamatsu) controlled by NIS-Elements Imaging Software. Images were obtained with a 60x 1.45NA objective lens with 200 nm z-step size over 12 μm depth. Laser intensity and exposure times were as follows: 100% and 300 ms for *cgh-1* smFISH (Quasar 670), 50% and 100 ms for endogenous PGL-1::GFP, and 5% and 100 ms for DAPI.

#### smFISH Analysis

mRNA spots were detected using the Big-FISH spot detection algorithm (version 0.6.2) ^107^, using a spot radius of 150 nm and a manually selected threshold of 27. The Big-FISH spot detection algorithm includes a step to detect bright regions with a potential cluster of spots and simulate a realistic number of spots within those regions. This decomposition was done by estimating the image background with a Gaussian filter and denoising the image, then aggregating pre-detected spots to build a reference spot. Gaussian parameters were fit to the reference spot and bright regions were detected as potential clusters. Then, as many Gaussians as possible were fit within the detected clusters. Clusters of mRNAs were defined as at least 6 mRNA spots within 350 nm distance. mRNAs within clusters were decomposed with the following Big-FISH parameters: alpha=0.75, beta=1.25, gamma=7. mRNA quantification was done using FISH-quant v2 and adapted Big-FISH Python script ^107^. For segmentation, embryos and germ cells were manually outlined using the polygon selection tool on Fiji^53^ using the Quasar670 and PGL-1::GFP channels as reference, respectively, and saved as ROIs. The embryos and germ cell ROIs were separately converted to binary images using the “ROIs to Label Image” function within the BIOP plugin. The binary images were used to generate “cell” (embryo) and “nuclei” (germ cell) boundaries. The numbers of mRNA molecules in germ and somatic cells were finally extracted using the analysis functions from Big-FISH. Damaged or inaccurately segmented embryos were excluded from the analyses.

#### Genetic Analyses

For embryonic lethality and brood size analyses, 1 L4 from each genotype was picked onto OP50-seeded NGM plates and grown at 16°C (or at 25°C for Figure 4B). After 8-12 hours of growth, P_0_ was transferred to a new plate and embryos laid on the original plate were manually counted, then repeated until P_0_ stopped laying progenies (typically 8-10 days). Surviving adult progenies from each plate were manually counted after 3-4 days post-P_0_ transfer. Embryonic lethality was calculated as the percentage of surviving adults over total embryos. For *let-7* analyses, 1 young adult was picked onto NGM plate as above, and the adult F1s and animals that died from vulval bursting during L4-to-adult transition were counted. For *lsy-6* analyses, 1 L4 was picked onto individual NGM plates and 100-200 adult F1s were analyzed under a 40x 0.75NA objective (Zeiss Axio Imager.M1 widefield microscope equipped with an AxioCam MRm camera) to analyze the number of *gcy-5p::GFP* ASER neuronal extensions. For imaging, adults were mounted on 2% agarose pads with 2 μL of 0.1% tetramisole (Sigma) in M9. For all analyses, at least 10 P_0_s (10 plates) were analyzed per genotype.

#### Data and Code Availability

The LC-MS/MS dataset was deposited in MassIVE: MSV000093990 and will be made public at the time of publication.

---

**Figure S1.**
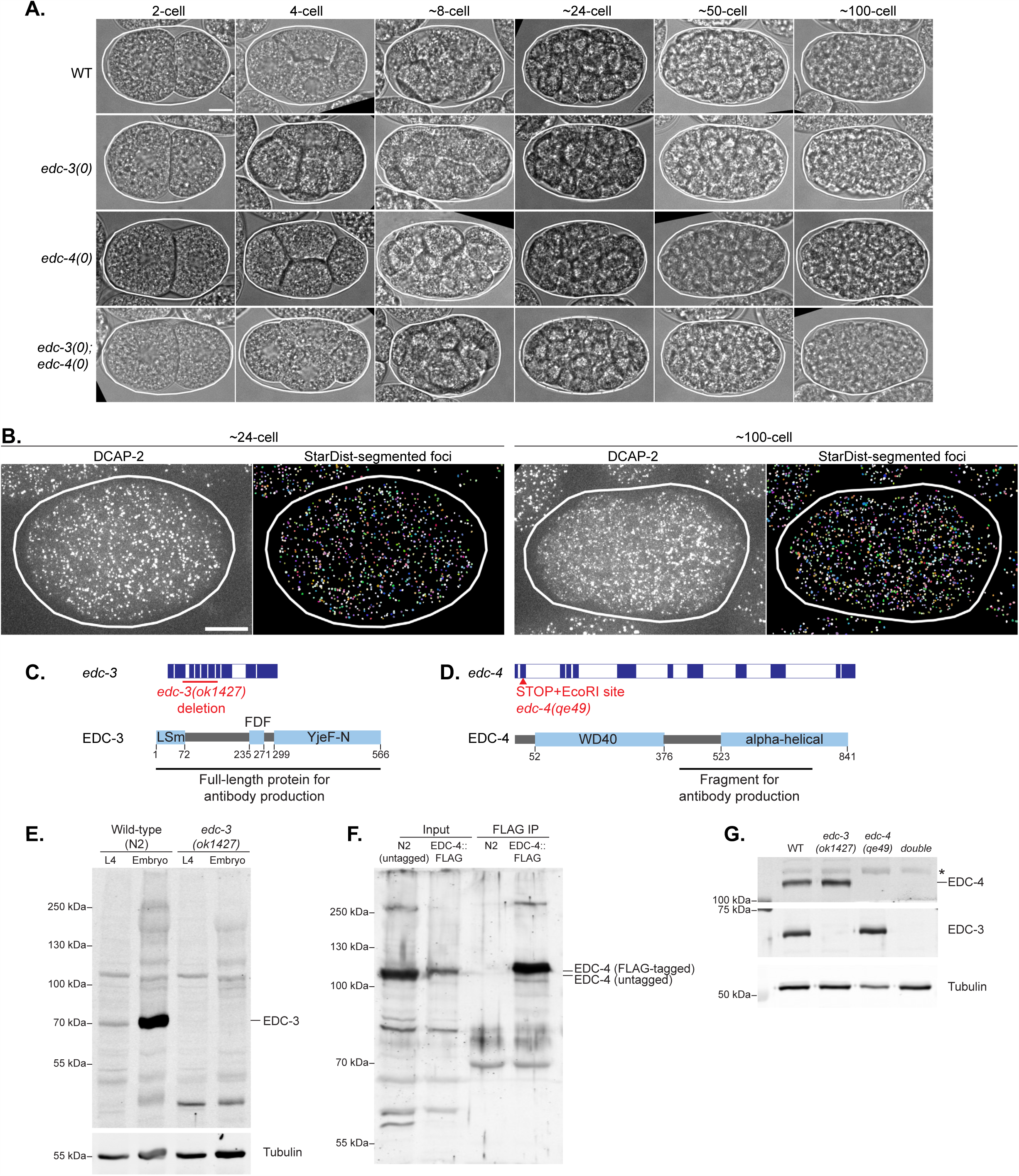
Refers to Figure 1. Corresponding brightfield photomicrographs and validation of EDC-3 and EDC-4 antibodies and alleles. (A) Corresponding brightfield images from Figure 1A. Scale bar is 10 µm. (B) Examples of DCAP-2 foci detected with the StarDist 2D plugin on Fiji. Scale bar is 10 µm. (C) Top: Schematic representation of *edc-3* genomic locus and location of the deletion in *edc-3(ok1427)* (i.e, *edc-3(0)*). Blue and white bars denote exons and introns, respectively. Bottom: Schematic representation of EDC-3 protein. (D) Top: Schematic representation of *edc-4* genomic locus and location of premature stop codon in *edc-4(qe49)* (i.e, *edc-4(0)*). Bottom: Schematic representation of EDC-4 protein. (E) Western blot validation of EDC-3 antibody on L4 and mixed-stage embryos lysates from WT (N2) or *edc-3(0)*. Tubulin serves as a loading control. (F) Western blot validation of EDC-4 antibody on embryonic lysates and FLAG immunoprecipitation of WT (N2) or a strain expressing *edc-4::3XFLAG*. (G) Western blot validation of EDC-3 and EDC-4 depletions in null alleles from embryonic lysates of WT (N2), *edc-3(0)*, *edc-4(0)*, or *edc-3(0); edc-4(0)* double mutant. Tubulin serves as a loading control. Star (*) indicates non-specific band.

**Figure S2.**
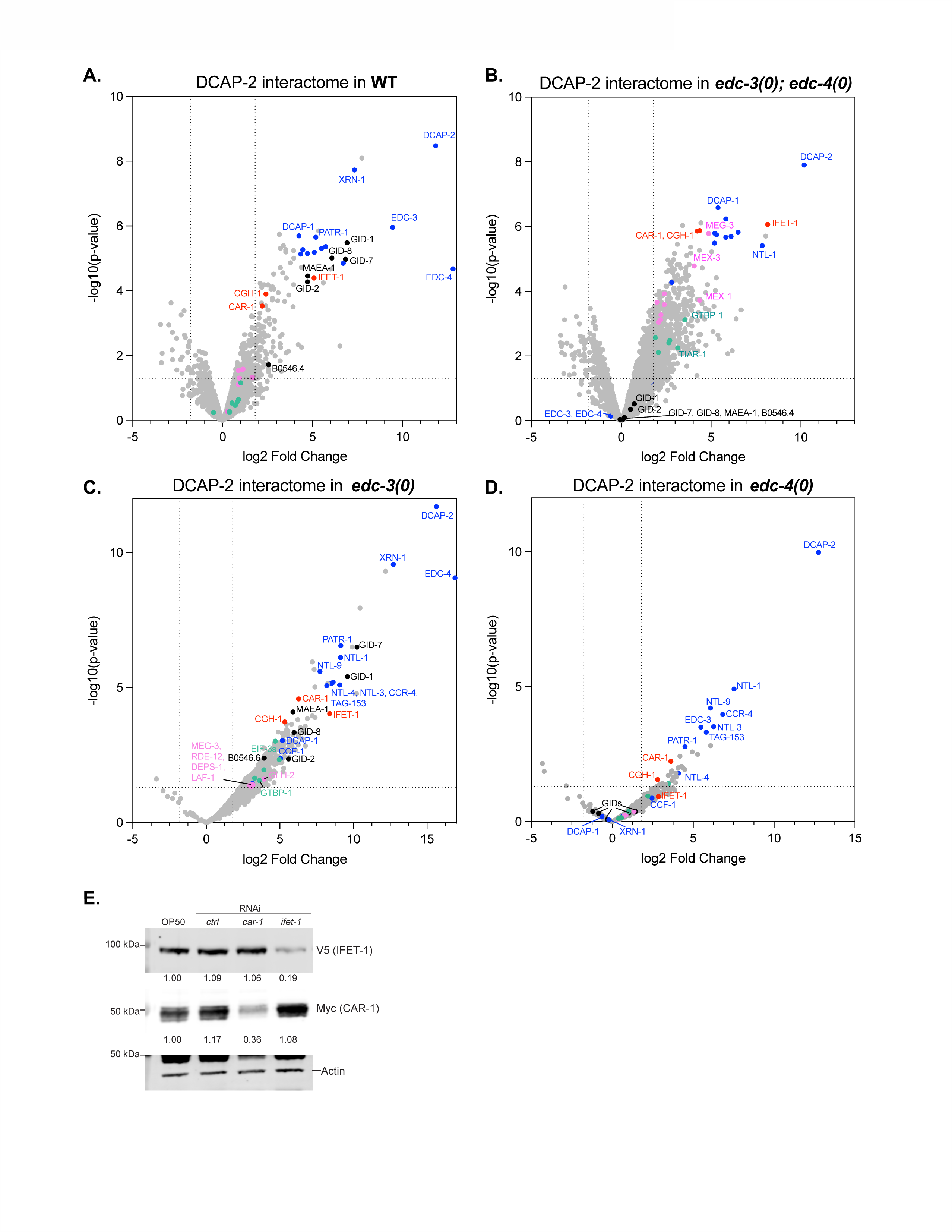
Refers to Figure 2. DCAP-2 interactome in WT and *edc-3/4* mutant embryos. (A-D) Volcano plots of proteins detected by LC-MS/MS of DCAP-2 immunoprecipitation (IP) eluates in (A) WT; (B) *edc-3(0); edc-4(0)* double mutant; (C) *edc-3(0)* single mutant; (D) *edc-4(0)*. P-values and log2 LFQ fold changes were computed using ProVision (see Methods). Proteins of interest were coloured as follows: Blue = P-body-enriched decapping/decay and deadenylation proteins; Red = germline-enriched decapping activators; Pink = Germ granule-enriched proteins; Green = Stress granule-enriched proteins. Black = GID complex subunits. Vertical dotted lines indicate log2(LFQ Fold Change) cut-off of 1.8. Horizontal dotted lines indicate -log10(p-value) = 1.3 (from Benjamini Hochberg FDR-corrected p-value cut-off of 0.05). Full list of proteins is listed in **Table S1**. Results were obtained from three biological and technical IP replicates. (E) Western blot validation of IFET-1 and CAR-1 depletion by RNAi on mixed-stage embryos expressing IFET-1::V5 and CAR-1::Myc. Actin served as a loading control. Numbers underneath each blot indicate fold change relative to OP50-fed group (no RNAi) and normalized to actin.

**Figure S3.**
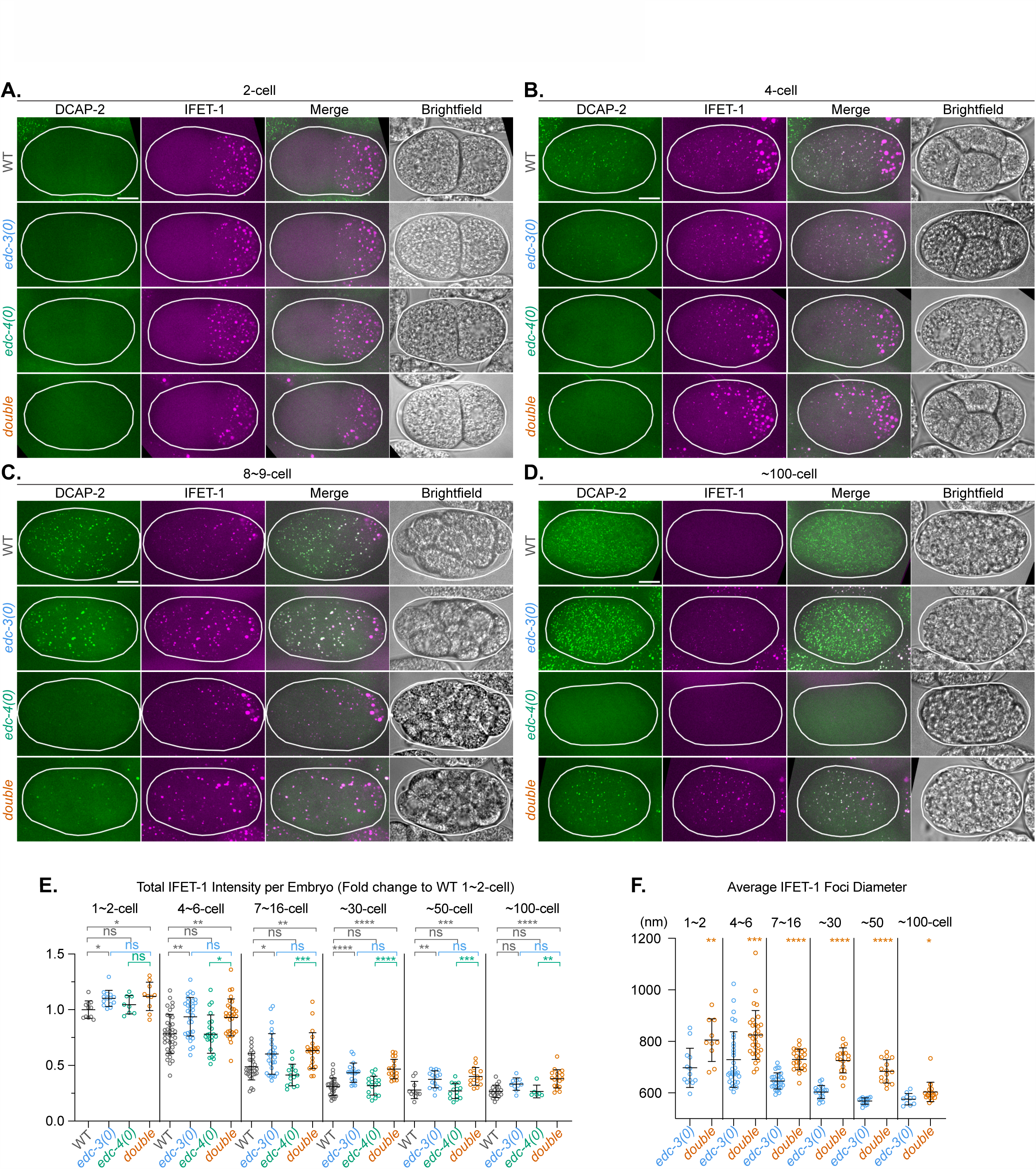
Refers to Figure 3. EDC-3 inhibits somatic accumulation of IFET-1. (A-D) Photomicrographs of live (A) 2-cell; (B) 4-cell; (C) 8-9-cell; (D) ∼100-cell embryos expressing IFET-1::TagRFP and DCAP-2::GFP in WT, *edc-3(0)*, *edc-4(0)*, and *edc-3(0); edc-4(0)* double mutant. Scale bars are 10 µM. (E) Total IFET-1 intensity per embryo. Represented as fold change of the mean intensity of all slices (sum projection), relative to WT at 4∼6-cell. Tukey’s multiple comparison test was conducted to determine statistical significance. (F) Average IFET-1 foci’s Feret’s diameter per embryo. Welch’s *t*-test was conducted to determine statistical significance. For (E) and (F), each data point represents the average from one embryo and is summarized as mean ± standard deviation. p-values: (*) =0.01-0.05, (**) =0.001-0.01, (***) =0.0001-0.001, (****) =<0.0001, p≥0.05 are insignificant (ns).

**Figure S4.**
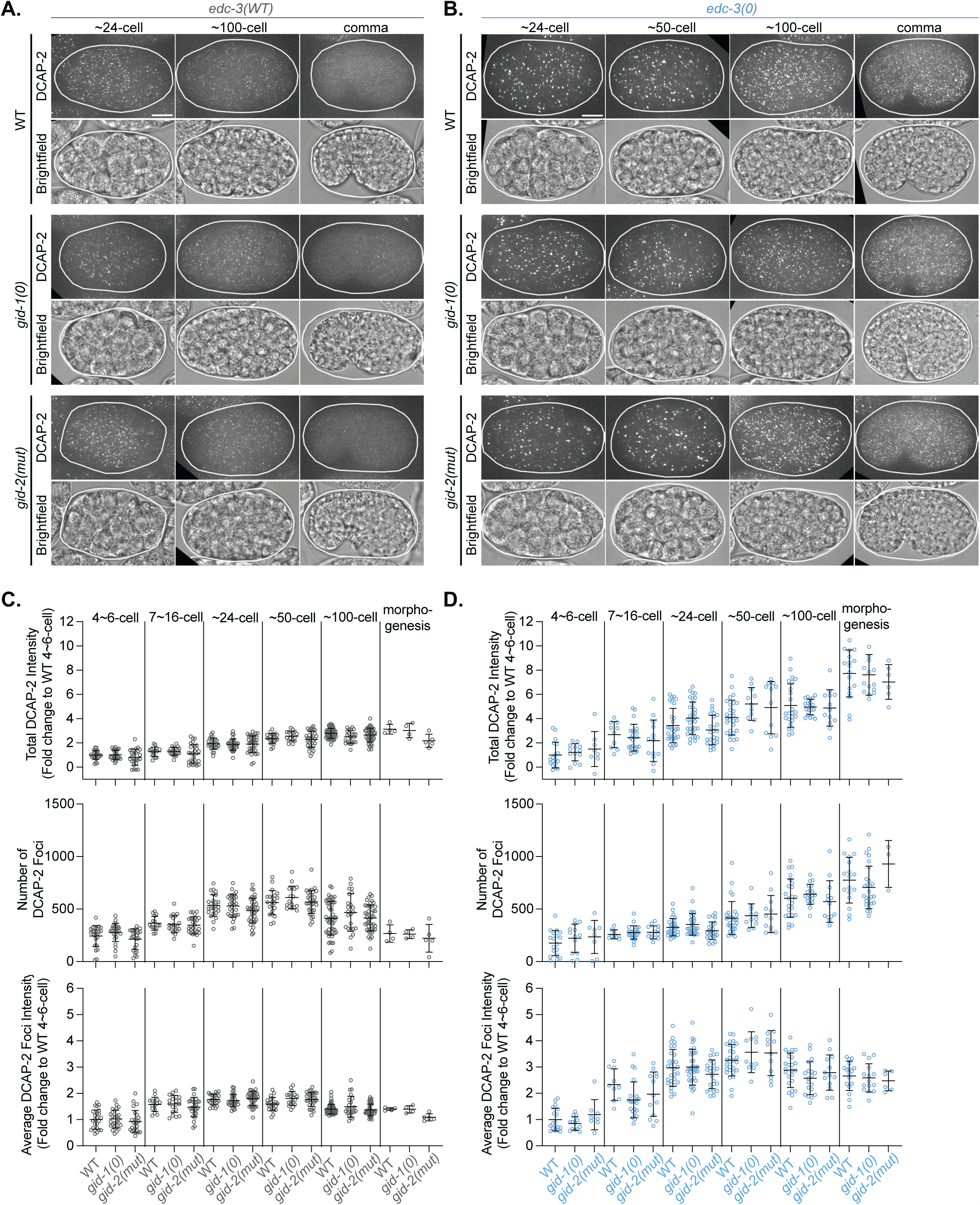
Refers to Figure 5. The GID complex is dispensable for DCAP-2 foci accumulation. (A-B) Corresponding DCAP-2::GFP and brightfield images from Figures 5C **and 5D**. Scale bars are 10 µm. (C-D) Total DCAP-2 intensity, number of DCAP-2 foci as detected with StarDist, and average DCAP-2 foci intensity in (C) WT or (D) *edc-3(0*). To note, WT in (B) and (D) refers to *edc-3(0); gid-1(WT); gid-2(WT)*. Results are representative of three technical and biological replicates wherein all genotypes were imaged in parallel. Each data point represents one embryo and data is summarized as mean ± standard deviation. Dunnet test was conducted to determine statistical significance relative to WT in each stage bin and differences were deemed insignificant (p≥0.05).

**Figure S5.**
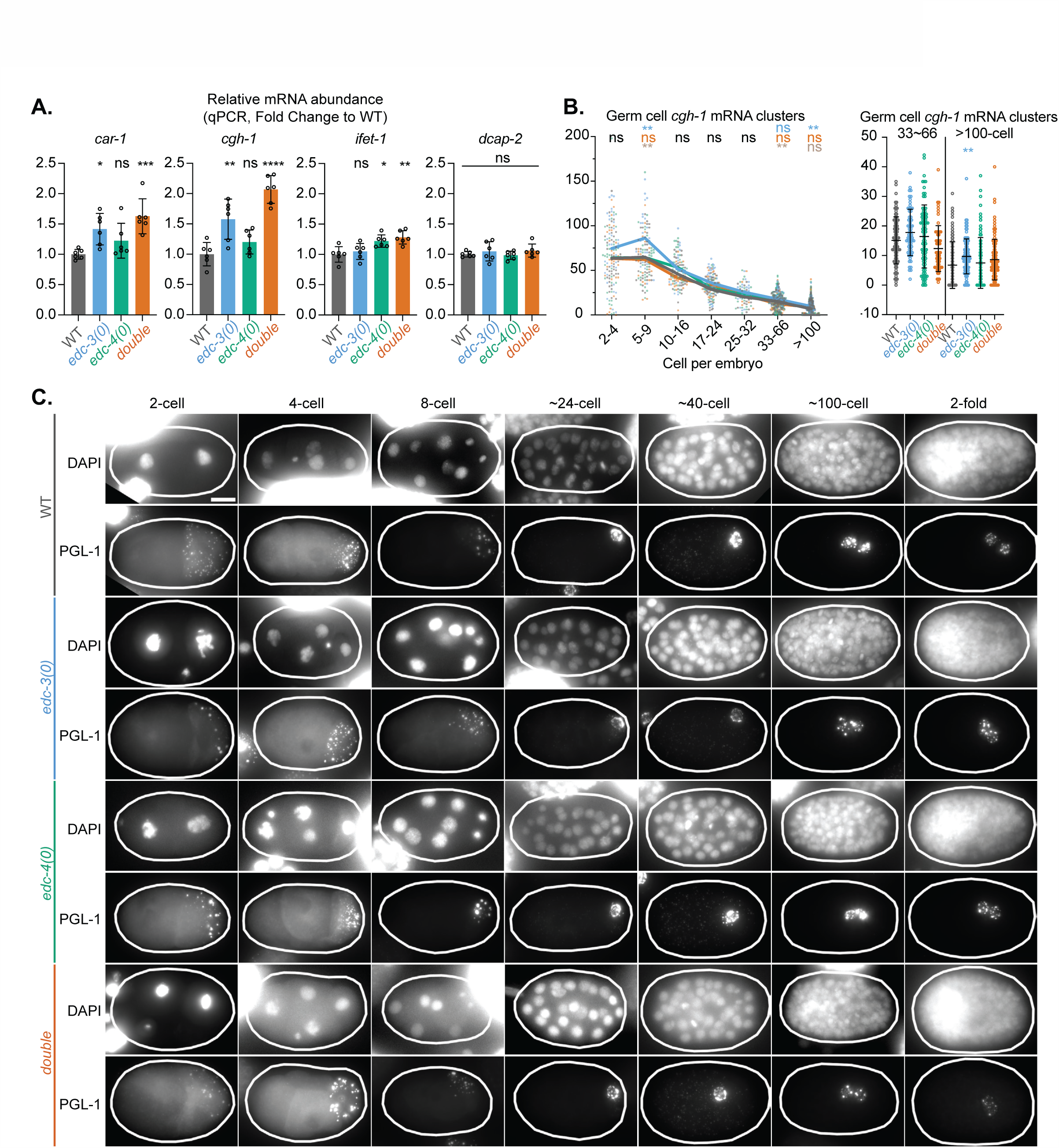
Refers to Figure 6. EDC-3 promotes the somatic and germline clearance of *cgh-1* mRNAs. (A) Relative abundance of *car-1*, *cgh-1*, *ifet-1*, and *dcap-2* mRNAs from mixed-stage embryos from WT, *edc-3(0)*, *edc-4(0)*, and *edc-3(0); edc-4(0)* double mutant, as measured by RT-qPCR and normalized to *act-1* mRNA. Fold change relative to WT is displayed as mean ± standard deviation from three biological replicates. Tukey’s multiple-comparisons test was conducted. p-values: (*) =0.01-0.05, (**) =0.001-0.01, (***) =0.0001-0.001, (****) =<0.0001, p≥0.05 are insignificant (ns). (B) Germ cell *cgh-1* mRNA cluster counts per embryo over development. (Right) Zoom of 34∼66 and >100-cell embryos to better visualize differences. (C) Corresponding DAPI and PGL-1::GFP images from Figure 6A. Scale bar is 10 µm.

